# Gene Networks and Pathways for Plasma Lipid Traits via Multi-tissue Multi-omics Systems Analysis

**DOI:** 10.1101/2020.02.28.970368

**Authors:** Montgomery Blencowe, In Sook Ahn, Zara Saleem, Helen Luk, Ingrid Cely, Ville-Petteri Mäkinen, Yuqi Zhao, Xia Yang

**Author notes:** **Correspondence should be addressed to:** Xia Yang, Ph.D., Department of Integrative Biology and Physiology, University of California Los Angeles, Los Angeles, CA 90095, Yuqi Zhao, Ph.D., Department of Integrative Biology and Physiology, University of California Los Angeles, Los Angeles, CA 90095. co-first author (Montgomery Blencowe, In Sook Ahn).

## Abstract

Genome-wide association studies (GWAS) have implicated ∼380 genetic loci for plasma lipid regulation. However, these loci only explain 17-27% of the trait variance and a comprehensive understanding of the molecular mechanisms has not been achieved. In this study, we utilized an integrative genomics approach leveraging diverse genomic data from human populations to investigate whether genetic variants associated with various plasma lipid traits, namely total cholesterol (TC), high and low density lipoprotein cholesterol (HDL and LDL), and triglycerides (TG), from GWAS were concentrated on specific parts of tissue-specific gene regulatory networks. In addition to the expected lipid metabolism pathways, gene subnetworks involved in ‘interferon signaling’, ‘autoimmune/immune activation’, ‘visual transduction’, and ‘protein catabolism’ were significantly associated with all lipid traits. Additionally, we detected trait-specific subnetworks, including cadherin-associated subnetworks for LDL, glutathione metabolism for HDL, valine, leucine and isoleucine biosynthesis for TC, and insulin signaling and complement pathways for TG. Finally, utilizing gene-gene relations revealed by tissue-specific gene regulatory networks, we detected both known (e.g. *APOH, APOA4*, and *ABCA1*) and novel (e.g. *F2* in adipose tissue) key regulator genes in these lipid-associated subnetworks. Knockdown of the *F2* gene (Coagulation Factor II, Thrombin) in 3T3-L1 and C3H10T1/2 adipocytes reduced gene expression of *Abcb11, Apoa5, Apof, Fabp1, Lipc*, and *Cd36*, reduced intracellular adipocyte lipid content, and increased extracellular lipid content, supporting a link between adipose thrombin and lipid regulation. Our results shed light on the complex mechanisms underlying lipid metabolism and highlight potential novel targets for lipid regulation and lipid-associated diseases.

## Introduction

Lipid metabolism is vital for organisms as it provides energy as well as essential materials such as membrane components and signaling molecules for basic cellular functions. Lipid dysregulation is closely related to many complex human diseases, such as atherosclerotic cardiovascular disease (1), Alzheimer’s disease (2, 3), type 2 diabetes (T2D) (4), and cancers (5). The notion of targeting lipid metabolism to treat human diseases has been reinforced by the fact that many disease-associated genes and drug targets (e.g., *HMGCR* as the target of statins and *PPARA* as the target of fibrates) are involved in lipid metabolic pathways (6-8).

Accumulating evidence supports that plasma lipids are complex phenotypes influenced by both environmental and genetic factors (9, 10). Heritability estimates for main plasma lipids are high (e.g. ∼70% for low density lipoprotein cholesterol [LDL] and ∼55% for high density lipoprotein cholesterol [HDL]) (11), indicating that DNA sequence variation plays an important role in explaining the inter-individual variability in plasma lipid levels. Indeed, genome-wide association studies (GWAS) have pinpointed a total of 386 genetic loci, captured in the form of single nucleotide polymorphisms (SNPs) associated with lipid phenotypes (12-16). For example, the most recent GWAS on lipid levels identified 118 loci that had not previously been associated with lipid levels in humans, revealing a daunting genetic complexity of blood lipid traits (16).

However, there are several critical issues that cannot be easily addressed by traditional GWAS analysis. First, even very large GWAS may lack statistical power to identify SNPs with small effect sizes and as a result the most significant loci only explain a limited proportion of the genetic heritability, for example, 17.2 – 27.1% for lipid traits (17). Second, the functional consequences of the genetic variants and the causal genes underlying the significant genetic loci are often unclear and await elucidation. To facilitate functional characterization of the genetic variants, genetics of gene expression studies (18, 19) and the ENCODE efforts (20) have documented tissue- or cell-specific expression quantitative trait loci (eQTLs) and functional elements of the human genome. These studies provide the much-needed bridge between genetic polymorphisms and their potential molecular targets. Third, the molecular mechanisms that transmit the genetic perturbations to complex traits or diseases, that is, the cascades of molecular events through which numerous genetic loci exert their effects on a given phenotype, remain elusive. Biological pathways that capture functionally related genes involved in molecular signaling cascades and metabolic reactions, and gene regulatory networks formed by regulators and their downstream genes can elucidate the functional organization of an organism and provide mechanistic insights (21). Indeed, various pathway- and network-based approaches to analyzing GWAS datasets have been developed (18, 22-24) and demonstrated to be powerful to capture both the missing heritability and the molecular mechanisms of many human diseases or quantitative phenotypes (18, 23, 25, 26). For these reasons, integrating genetic signals of blood lipids with multi-tissue multi-omics datasets that carry important functional information may provide a better understanding of the molecular mechanisms responsible for lipid regulation as well as the associated human diseases.

In this study, we apply an integrative genomics framework to identify important regulatory genes, biological pathways, and gene subnetworks in relevant tissues that contribute to the regulation of four critical blood lipid traits, namely TC, HDL, LDL, and TG. We combine the GWAS results from the Global Lipids Genetics Consortium (GLGC) with functional genomics data from a number of tissue-specific eQTLs and the ENCODE project, and gene-gene relationship information from biological pathways and data-driven gene network studies. The integrative framework is comprised of four main parts (**Figure 1**): 1) Marker Set Enrichment Analysis (MSEA) where GWAS, functional genome, and pathways or co-regulated genes are integrated to identify lipid-related functional units of genes, 2) merging and trimming of identified lipid gene sets, 3) key driver analysis (KDA) to pinpoint important regulatory genes by further integrating gene regulatory networks, and 4) validation of key regulators using genetic perturbation experiments and *in silico* evidence. This integrated systems biology approach enables us to derive a comprehensive view of the complex and novel mechanisms underlying plasma lipid metabolism.

**Figure 1.**
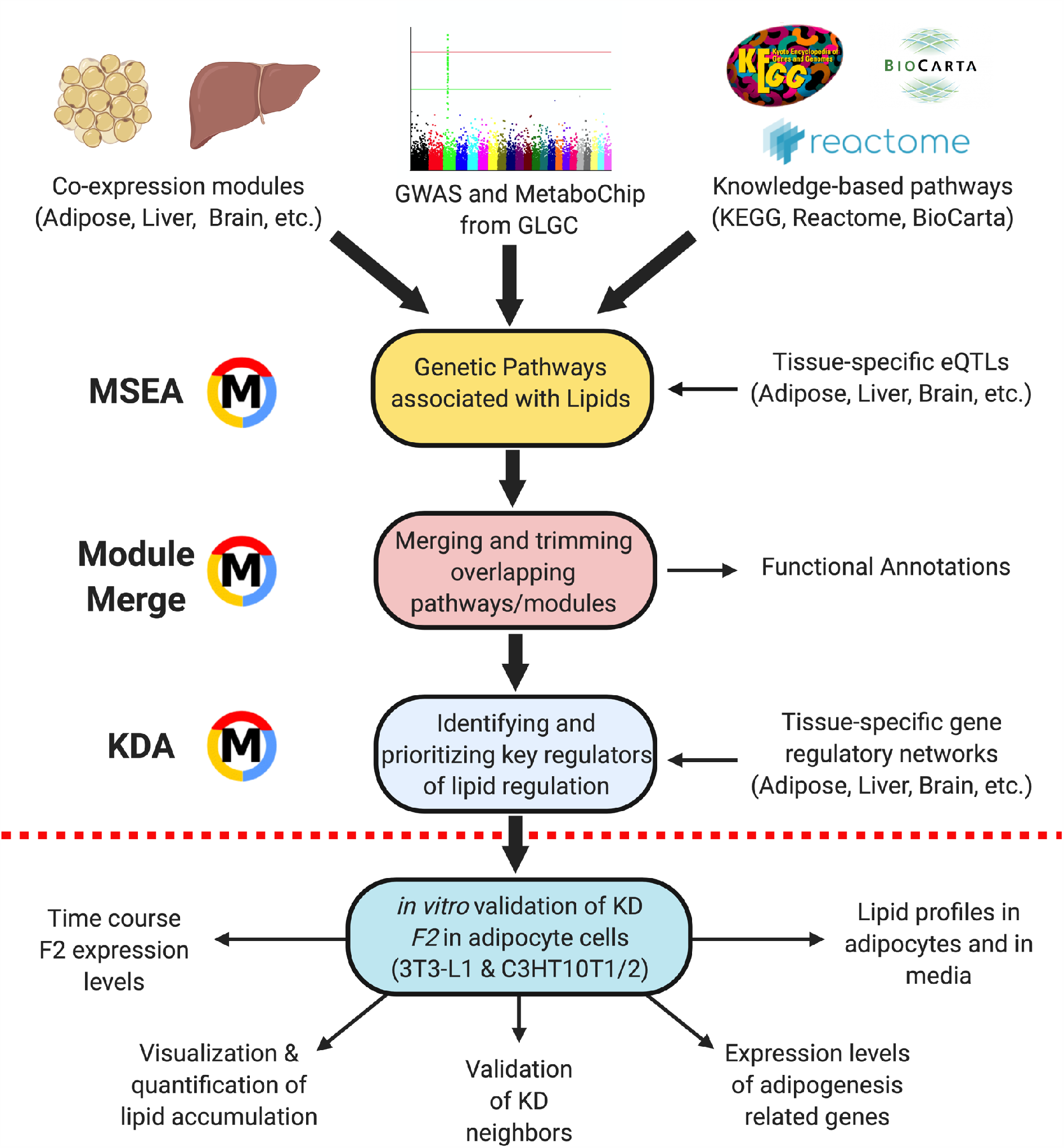
Overall design of the study. The statistical framework can be divided into four main parts, including Marker Set Enrichment Analysis (MSEA), merging and trimming of gene sets, Key Driver Analysis (KDA), and validation of the key regulators using *in vitro* testing.

## Materials and Methods

### GWAS of lipid traits

The experimental design, genotyping, and association analyses of HDL, LDL, TC, and TG were described previously (12). The dataset used in this study is comprised of > 100,000 individuals of European descent (sample size 100,184 for TC, 95,454 for LDLC, 99,900 for HDLC and 96,598 for TG), ascertained in the United States, Europe, or Australia. More than 906,600 SNPs were genotyped using Affymetrix Genome-Wide Human SNP Array 6.0. Imputation was further carried out to obtain information for up to 2.6 million SNPs using the HapMap CEU (Utah residents with ancestry from northern and western Europe) panel. SNPs with minor allele frequency (MAF) < 1% were removed. Finally, a total of ∼ 2.6 million SNPs tested for association with each of the four lipid traits were used in our study.

### Genetic association study of lipid traits using MetaboChip

The experimental design, genotyping, and association analyses of the lipid MetaboChip study were described previously (27). The study examined subjects of European ancestry, including 93,982 individuals from 37 studies genotyped with the MetaboChip array, comprised of 196,710 SNPs representing candidate loci for cardiometabolic diseases. There was limited overlap between the individuals involved in GWAS and those in MetaboChip.

### Knowledge-based biological pathways

We included canonical pathways from the Reactome (version 45), Biocarta, and the Kyoto Encyclopedia of Genes and Genomes (KEGG) databases (28, 29). In addition to the curated pathways, we constructed four positive control pathways based on known lipid-associated loci (p < 5.0 × 10^−8^) and candidate genes from the GWAS Catalog (30). These gene sets were based on 4, 11, 13, and 13 studies for TC, TG, LDL, and HDL, respectively (full lists of genes in each positive control sets are in **Supplemental Table S1**) and serve as positive controls to validate our computational method.

### Data-driven modules of co-expressed genes

Beside the canonical pathways, we used co-expression modules that were derived from a collection of genomics studies (**Supplement Table S2**) of liver, adipose tissue, aortic endothelial cells (HAEC), brain, blood, kidney, and muscle (31-40). A total of 2706 co-expression modules were used in this study. Although liver and adipose tissue are likely the most important tissues for lipid regulation, we included the other tissue networks to confirm whether known tissue types for lipids could be objectively detected and whether any additional tissue types are also important for lipids.

### Mapping SNPs to genes

Three different mapping methods were used in this study to link SNPs to their potential target genes.

#### Chromosomal distance-based mapping

First, we used a standard distance-based approach where a SNP was mapped to a gene if within 50 kb of the respective gene region. The use of ± 50 kb to define gene boundaries is commonly used in GWAS.

#### eQTL-based mapping

The expression levels of genes can be seen also as quantitative traits in GWAS. Hence, it is possible to determine eQTLs and the expression SNPs (eSNPs) within the eQTLs that provide a functionally motivated mapping from SNPs to genes. Moreover, the eSNPs within the eQTL are specific to the tissue where the gene expression was measured and can therefore provide mechanistic clues regarding the tissue of action when intersected with lipid-associated SNPs. Results from eQTL studies in human adipose tissue, liver, brain, blood, and HAEC were used in this study (31, 33, 34, 41-49). We included both *cis*-eSNPs (within 1 Mb distance from gene region) and *trans*-eSNPs (beyond 1 Mb from gene region), at a false discovery rate < 10%.

#### ENCODE-based mapping

In addition to eQTLs and distance-based SNP-gene mapping approaches, we integrated functional data sets from the Regulome database (20), which annotates SNPs in regulatory elements in the *Homo sapiens* genome based on the results from the ENCODE studies (50).

#### Nine unique combinations of SNP-gene mapping

Using the above three mapping approaches, we derived nine unique sets of SNP-gene mapping. These are: eSNP adipose, eSNP liver, eSNP blood, eSNP brain, eSNP HAEC, eSNP all (i.e., combining all the tissue-specific eSNPs above), Distance (chromosomal distance-based mapping), Regulome (ENCODE-based mapping), and Combined (combining all the above methods).

### Removal of SNPs in linkage disequilibrium

We observed a high degree of linkage disequilibrium (LD) in the eQTL, Regulome, and distance-based SNPs, and this LD structure may cause artifacts and biases in the downstream analysis. For this reason, we devised an algorithm to remove SNPs in LD while preferentially keeping those with a strong statistical association with lipid traits. Technical details are available in Supplementary Methods. We chose a LD cutoff (R^2^ < 0.5) to remove redundant SNPs in high LD.

### Marker Set Enrichment Analysis (MSEA)

We applied a modified MSEA method (24, 51) to find pathways/co-expressed modules associated with lipid traits (**Supplemental Methods**). False discovery rates (FDR) were estimated with the method by Benjamini and Hochberg (52). Pathways or co-expression modules with FDR < 10% were considered statistically significant. MSEA were applied to both the GLGC GWAS dataset and the MetaboChip dataset. The combined FDR from these two datasets was expected to be < 1% (10% * 10% = 1%).

### Comparison between MSEA and other computational method

To ensure that the pathway results from MSEA are reproducible, we used the improved gene-set-enrichment analysis approach (iGSEA) (53). In the iGSEA analysis, we generated gene sets using the same canonical pathways and co-expression modules in MSEA. The SNPs were mapped to genes using the default settings of iGSEA. For each given gene set, significance proportion-based enrichment score was calculated to estimate the enrichment of genotype–phenotype association. Then, iGSEA performed label permutations to calculate nominal P-values to assess the significance of the pathway-based enrichment score and FDR to correct multiple testing, with FDR < 0.25 (default setting) regarded as significant pathways. Considering that MSEA and iGSEA were independent, the combined FDR from these two methods of analysis was expected to be < 5% (10% × 25% = 2.5%).

### Construction of independent supersets and confirmation of lipid association

Because the pathways or co-expression modules were collected from multiple sources, there were overlapping or nested structures among the gene sets. To make the results more meaningful, we constructed relatively independent supersets that captured the core genes from groups of redundant pathways and co-expression modules (**Supplemental Methods**). After merging, we annotated each superset based on function enrichment analysis of the known pathways from the Gene Ontology and KEGG databases (P < 0.05 in Fisher’s exact test after Bonferroni correction). The supersets were given a second round of MSEA to confirm their significant associated with lipids using P < 0.05 after Bonferroni correction as the cutoff.

### Key driver analysis (KDA)

We adopted a previously developed KDA algorithm (54-56) of gene-gene interaction networks to the lipid-associated supersets in order to identify the key regulatory genes (**Figure 1**). In the study, we included Bayesian gene regulatory networks from diverse tissues, including adipose tissue, liver, blood, brain, kidney and muscle (31-39). A key driver was defined as a gene that is directionally connected to a large number of genes from a lipid superset, compared to the expected number for a randomly selected gene within the Bayesian network (details in **Supplemental Methods**). The MSEA, merging, and KDA were performed using R.

### Enrichment analysis of lipid-associated subnetworks in human complex diseases

We collected disease susceptibility genes from GWAS Catalog with GWAS P<10E-5 for four human complex diseases, including cardiovascular diseases (‘myocardial infarction’, ‘myocardial infarction (early onset)’, ‘coronary artery calcification’, and ‘coronary heart disease’), Alzheimer’s disease, type 2 diabetes, and cancer (‘colon cancer’, ‘breast cancer’, ‘pancreas cancer’, ‘prostate cancer’, and ‘chronic lymphocytic leukemia’). Fisher’s exact test was used to explore the enrichment of genes in the lipid-associated subnetworks in the disease gene sets. Bonferroni-corrected p < 0.05 was considered significant.

### Validation of *F2* in adipocyte functions via *F2* siRNA transfection in 3T3-L1 and C3H10T1/2 adipocyte cell lines

The mouse preadipocytes 3T3-L1 and C3H10T1/2 cells were obtained from ATCC and maintained and differentiated to adipocytes according to the manufacturer’s instruction. For knockdown experiments, 3 predesigned siRNAs targeting *F2* gene (sequences in **Supplemental Table S3**; GenePharma, Paramount, CA) were tested and the most effective one was selected for the experiment (**Supplemental Figure S1**). We first measured *F2* expression during adipocyte differentiation and found increased *F2* expression on days 8-10 in 3T3-L1 and days 6-10 in C3H10T1/2 during differentiation, which helped inform on the timing of siRNA transfection in these cell lines. 3T3-L1 adipocytes were transfected with 50 nM of *F2* siRNA using Lipofectamin 2000 on day 7 (D7) of differentiation, a day before *F2* increase. Followed by 72 hrs of siRNA treatment, adipocytes were processed for Oil red O staining of lipids and Real-time qPCR for select genes. C3H10T1/2 adipocytes were transfected with 50 nM of *F2* siRNA using Lipofectamin 2000 on day 5 (D5) and day 7 (D7), and adipocytes were processed on day 9 (D9) for Oil red O staining of lipids, Real-time qPCR for select genes, and quantitative lipid assays. As control, 50 nM of scrambled siRNA (GenePharma, Paramount, CA) was transfected at the same time points as the *F2* siRNA in the two cell lines. To determine changes in lipid accumulation, adipocytes were stained by Oil red O stain solution. After obtaining images, Oil red O was eluted in isopropyl alcohol and optical density (OD) values were measured at 490 nm.

### RNA extraction and Real-time qPCR

Total RNA was extracted from the adipocytes (Zymo Research, Irvine, CA), and RNA was reverse transcribed using cDNA Reverse Transcription Kit (Thermo Scientific, Madison, WI, USA), Real-time qPCR for select network and non-network genes was performed using the primers shown in **Supplemental Table S3**. Each reaction mixture (20 ul) is composed of PowerUp SYBR Green Master Mix (Applied Biosystems), 0.5 uM each primer, and cDNA (150 ng for *F2* gene, 20-50 ng for the other genes), Each sample was tested in duplicate under the following amplification conditions: 95°C for 2 min, and then 40 cycles of 95°C for 1 s and 60°C for 30 s in QuantStudio 3 Real-Time PCR System (Applied Biosystems, Foster City, CA, USA). PCR primers were designed using the Primer-BLAST tool available from the NCBI web site (57). Melt curve was checked to confirm the specificity of the amplified product. Relative quantification was calculated using the 2 ^ (-ΔΔ CT) method (58). Beta actin was used as an endogenous control gene to evaluate the gene expression levels. All data are presented as the mean ± s.e.m of n = 4/group. Statistical significance was determined by Two-tailed Student’s t test and values were considered statistically significant at *P* < 0.05.

### Extraction and quantification of lipids in cells and media

Lipids were extracted from C3H10T1/2 cells and culture media using the Folch method (59) with minor modifications. Briefly, whole culture medium (1 mL) from each well of 12-well plate was collected in a separate tube. Cells were washed with phosphate buffered saline (PBS), and collected in 1 mL PBS and homogenized. Media or cell homogenate was mixed in 5 ml of chloroform: methanol (2:1, vol/vol) by shaking vigorously several times, and centrifuged at 2,500 × g for 15 min. Bottom organic layer was transferred to a new glass tube. The remaining aqueous phase and interphase including soluble protein were mixed with 5 mL chloroform by vigorous shaking, followed by centrifugation at 2,500 × g for 15 min. Bottom organic layer was combined with the first collected organic layer. The combined organic phase was evaporated using nitrogen, and then the dried lipids were resuspended in 0.5 % Triton X-100 in water. Samples were stored in - 80°C until lipid analysis. Triglyceride (TG), total cholesterol (TC), unesterified cholesterol (UC), and phospholipid (PL) levels in lipid extractions from cells and from culture media were measured separately using a colorimetric assay at the UCLA GTM Mouse Transfer Core (60). Intracellular lipids were normalized to the cellular protein amount measured by BCA protein assay kit (Pierce, Rockford, IL, USA). Extracellular lipids are presented as lipid quantity in 1 mL of collected media.

## Results

### Identification of pathways and gene co-expression modules associated with lipid traits

To asses biological pathway enrichment for the four lipid traits with GLGC GWAS, we curated a total of 4532 gene sets including 2705 tissue-specific co-expression modules (i.e., highly co-regulated genes based on tissue gene expression data) and 1827 canonical pathways from Reactome, Biocarta and the Kyoto Encyclopedia of Genes and Genomes (KEGG). These gene sets were constructed as data- and knowledge-driven functional units of genes. Four predefined positive control gene sets for HDL, LDL, TC, and TG were also created based on candidate genes curated from the GWAS catalog (61). To map potential functional SNPs to genes in each gene set, tissue-specific eQTLs, ENCODE functional genomics information, and chromosomal distance-based mapping were used (details in Methods). Tissue-specific eQTL sets were obtained from the GTEx database from studies on human adipose tissue, liver, brain, blood, and human aortic endothelial cells (HAEC), and a total of nine SNP-gene mapping methods were created. The liver and adipose tissues have established roles in lipid regulation, whereas the other tissues are included for comparison.

Integrating the datasets mentioned above using MSEA, we identified 65, 86, 90, and 92 gene sets whose functional genetic polymorphisms showed significant association with HDL, LDL, TC, and TG, respectively, in GLGC GWAS (FDR < 10%; **Supplemental Table S4)**. The predefined positive controls for the four lipid traits were among the top signals for their corresponding traits (**Table 1**), indicating that our MSEA method is sensitive in detecting true lipid trait-associated processes. Compared with other tissues, more pathways were captured when using liver and adipose eSNPs to map GWAS SNPs to genes (**Supplemental Table S4)**. For example, 56 out of the 86 LDL-associated pathways were found when liver and adipose eSNPs were used in our analysis. These results confirmed the general notion that liver and adipose tissue play critical roles in regulating plasma lipids, leading us to focus the bulk of our analysis on these two tissues, with the remaining tissues serving as a supplement.

**Table 1.** Common pathways shared by the four lipid traits in SNP set enrichment analysis.

| Categories | Descriptions | Traits* |  |  |  | MetaboChip | iGSEA |
| --- | --- | --- | --- | --- | --- | --- | --- |
|  |  | HDL | LDL | TC | TG |  |  |
| Positive Controls | Positive control gene set for TG | 1,2,3,5,6,7,8,9 | 2,3,5,6,7,8,9 | 2,3,5,6,7,8,9 | 1,2,3,5,6,7,8,9 | YES | YES |
|  | Positive control gene set for LDL | 5,6,7,8,9 | 1,2,3,4,5,6,7,8,9 | 1,2,3,4,5,6,7,8,9 | 1,2,3,5,6,7,8,9 | YES | YES |
|  | Positive control gene set for TC | 3,5,6,7,8,9 | 1,2,3,4,5,6,7,8,9 | 1,2,3,4,5,6,7,8,9 | 1,2,3,5,6,7,8,9 | YES | YES |
|  | Positive control gene set for HDL | 1,2,3,4,5,6,7,8,9 | 2,6,7,8,9 | 2,5,6,7,8,9 | 1,2,5,6,7,8,9 | YES | YES |
| Lipid metabolism | Lipoprotein metabolism | 1,2,5,6,7,8,9 | 5,6,7,8,9 | 5,6,7,8,9 | 5,6,7,8,9 | YES | YES |
|  | Chylomicron-mediated lipid transport | 5,6,7,8,9 | 7,8,9 | 5,6,7,8,9 | 5,6,7,8,9 | YES | YES |
|  | LDL-mediated lipid transport | 6,7,9 | 6,7,9 | 6,7,9 | 6,7,9 | NO | YES |
|  | HDL-mediated lipid transport | 1,2,5,6,7,8,9 | 5,7,8,9 | 5,7,8,9 | 5,7,8,9 | YES | YES |
| Protein catabolism | ER-Phagosome pathway | 1,5,8,9 | 1,3,5,6,8,9 | 1,2,3,5,6,8,9 | 1,3,5,6,8,9 | YES | YES |
|  | Antigen processing and presentation | 5,9 | 1,2,3,5,6,7,8,9 | 1,2,3,5,6,7,8,9 | 1,2,3,5,6,7,8,9 | YES | YES |
| Interferon Signaling | Interferon Signaling | 7,9 | 1,3,5,6,8,9 | 1,2,3,5,6,8,9 | 1,3,5,8 | YES | YES |
| Autoimmune/Immune activation | Type I diabetes mellitus | 1,5 | 1,2,3,5,6,7,8,9 | 1,2,3,5,6,7,8,9 | 1,2,3,5,6,7,8,9 | YES | YES |
|  | Scavenging by Class B Receptors | 6,7,8,9 | 7,9 | 7,9 | 7,9 | NO | YES |
|  | Asthma | 6 | 1,3,5,6,7,8,9 | 1,2,3,5,6,7,8,9 | 1,2,3,5,6,7,8,9 | YES | YES |
|  | IL 5 Signaling Pathway | 5 | 1,5,6,8,9 | 1,5,6,8,9 | 5,6,8 | NO | NO |
|  | Th1/Th2 Differentiation | 3 | 1,3,5,6,8 | 1,3,5,6,8,9 | 1,3,5,6,8 | NO | YES |
|  | Natural killer cell mediated cytotoxicity | 5 | 1,3,5 | 1,3,5,6,9 | 1,3,5 | YES | YES |
|  | HLA genes | 1,3,5,6,7,8,9 | 1,2,3,5,6,7,8,9 | 1,2,3,5,6,7,8,9 | 1,2,3,5,6,7,8,9 | YES | YES |
|  | Cell adhesion molecules (CAMs) | 5 | 1,2,3,5,6,7,8,9 | 1,2,3,5,6,7,8,9 | 1,3,5,6,8,9 | YES | NO |
|  | Autoimmune thyroid disease | 1,3,5,6,8,9 | 1,2,3,5,6,7,8,9 | 1,2,3,5,6,7,8,9 | 1,2,3,5,6,7,8,9 | YES | YES |
| Visual transduction | Diseases associated with visual transduction | 7 | 7,8,9 | 7,8,9 | 7,9 | YES | YES |
|  | Visual phototransduction | 7 | 7,8,9 | 7,8,9 | 7,9 | YES | YES |

Note: \*: The trait columns represent in which methods the MSEA of the pathways is significant with FDR
< 10%. Number 1 to 9 represent: adipose eSNP (1), blood eSNP (2), brain eSNP (3), human aortic endothelial cells (HAEC) eSNP (4), liver eSNP (5), all eSNP (6), Distance (7), Regulome (8), and Combined (9), respectively. The MetaboChip and iGSEA columns tell whether the gene set can also be detected as statistically significant in the analysis.

Among the significant gene sets, 39 were shared across the four lipid traits. These gene sets represented the expected lipid metabolic pathways as well as those that are less known to be associated with lipids, such as ‘antigen processing and presentation’, ‘cell adhesion molecules (CAMs)’, ‘visual phototransduction’, and ‘IL-5 signaling pathway’ (summary in **Table 1**; details in **Supplemental Table S4)**. We broadly classified the common gene sets detected into ‘positive controls’, ‘lipid metabolism’, ‘interferon signaling’, ‘autoimmune/immune activation’, ‘visual transduction’, and ‘protein catabolism’ (**Table 1**).

Beside the common gene sets described above, we also detected 18, 5, 6, and 17 trait-specific pathways/modules for HDL, LDL, TC, and TG, respectively (**Table 2**; **Supplement Table S4**), suggesting trait-specific regulatory mechanisms. Among the 18 pathways for HDL were ‘cation-coupled chloride transporters’, ‘glycerolipid metabolism’ and ‘negative regulators of RIG-I/MDA5 signaling’ across analyses using different tissue eSNP mapping methods, ‘alcohol metabolism’ from brain-based analysis, ‘packaging of telomere ends’ in adipose tissue, ‘glutathione metabolism’ in liver, and ‘cobalamin metabolism’ and ‘taurine and hypotaurine metabolism’ in both adipose and liver-based analyses. LDL-specific pathways included the ‘platelet sensitization by LDL’ pathway and a liver co-expression module related to cadherin. TC-specific pathways included ‘valine, leucine and isoleucine biosynthesis’ across tissues and ‘wound healing’ in the brain-based analysis. When looking at the TG-specific pathways, gene sets associated with ‘cellular junctions’ were consistent across tissues whereas ‘insulin signaling’ and complement pathways were exclusively seen in adipose tissue-based analysis.

**Table 2.**
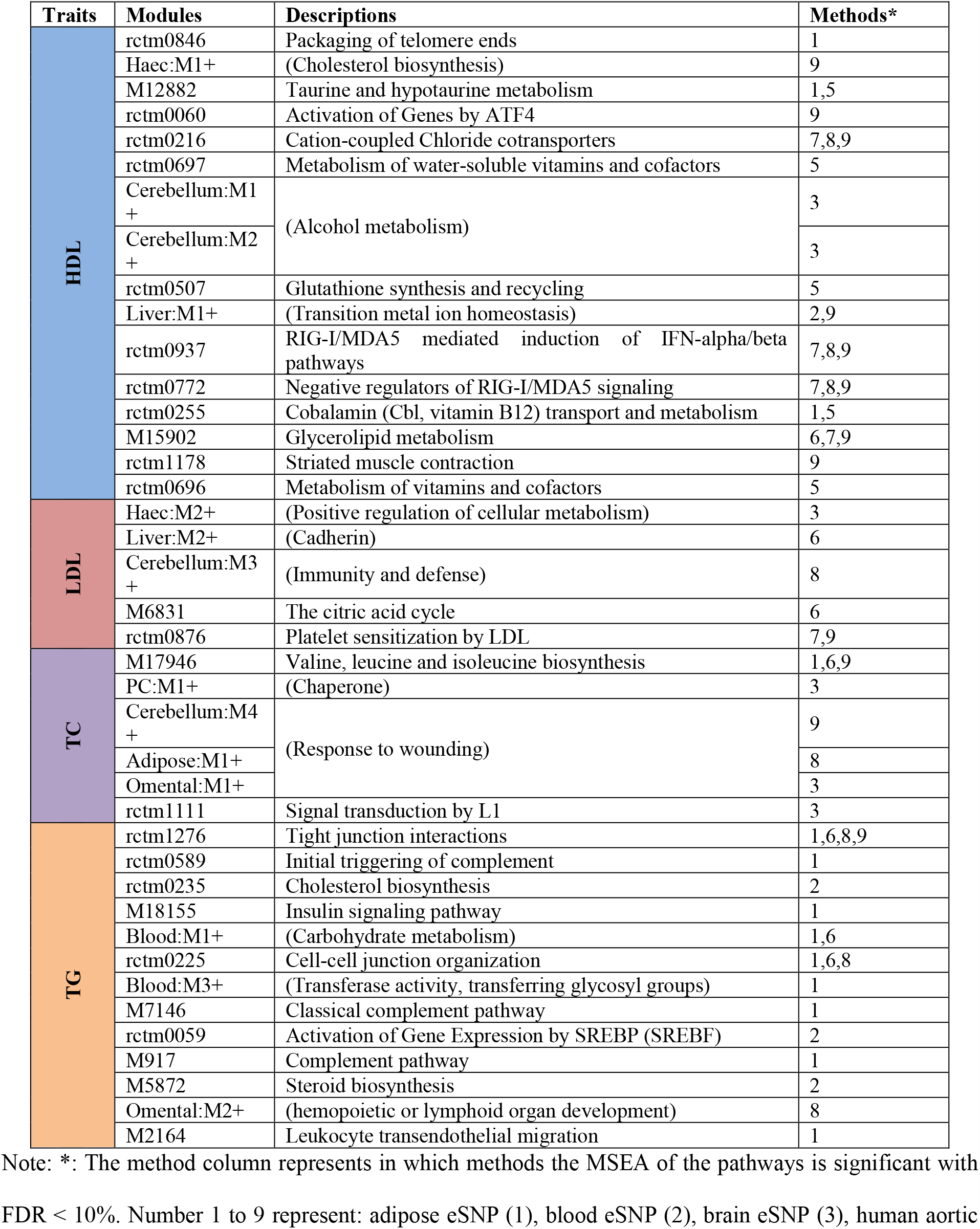
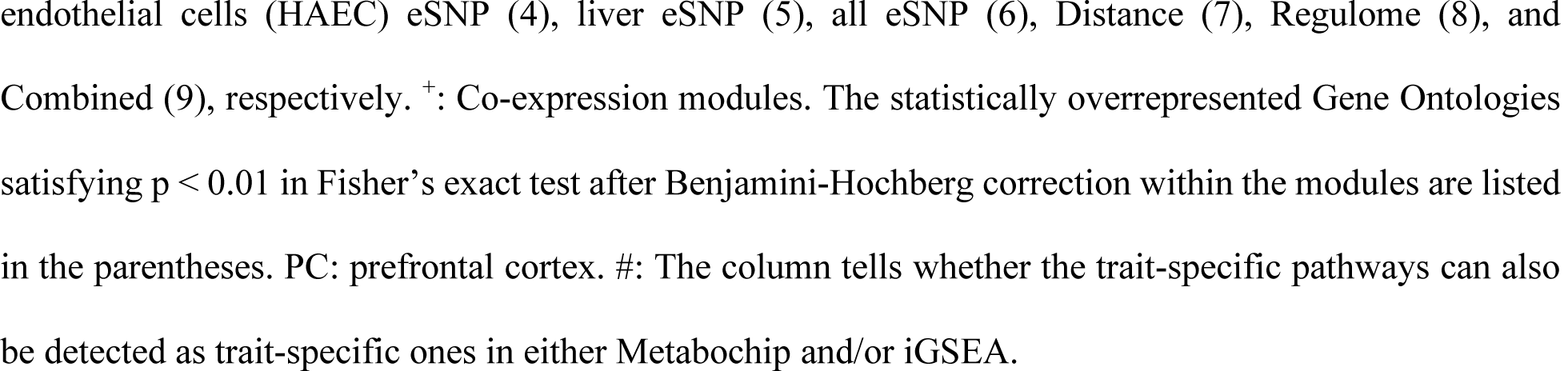
Trait-specific pathways identified in the SNP set enrichment analysis for four lipid traits.

| Traits | Modules | Descriptions | Methods* |
| --- | --- | --- | --- |
| HDL | rctm0846 | Packaging of telomere ends | 1 |
|  | Haec:M1+ | (Cholesterol biosynthesis) | 9 |
|  | M12882 | Taurine and hypotaurine metabolism | 1,5 |
|  | rctm0060 | Activation of Genes by ATF4 | 9 |
|  | rctm0216 | Cation-coupled Chloride cotransporters | 7,8,9 |
|  | rctm0697 | Metabolism of water-soluble vitamins and cofactors | 5 |
|  | Cerebellum:M1+ | (Alcohol metabolism) | 3 |
|  | Cerebellum:M2+ |  | 3 |
|  | rctm0507 | Glutathione synthesis and recycling | 5 |
|  | Liver:M1+ | (Transition metal ion homeostasis) | 2,9 |
|  | rctm0937 | RIG-I/MDA5 mediated induction of IFN-alpha/beta pathways | 7,8,9 |
|  | rctm0772 | Negative regulators of RIG-I/MDA5 signaling | 7,8,9 |
|  | rctm0255 | Cobalamin (Cbl, vitamin B12) transport and metabolism | 1,5 |
|  | M15902 | Glycerolipid metabolism | 6,7,9 |
|  | rctm1178 | Striated muscle contraction | 9 |
|  | rctm0696 | Metabolism of vitamins and cofactors | 5 |
| LDL | Haec:M2+ | (Positive regulation of cellular metabolism) | 3 |
|  | Liver:M2+ | (Cadherin) | 6 |
|  | Cerebellum:M3+ | (Immunity and defense) | 8 |
|  | M6831 | The citric acid cycle | 6 |
|  | rctm0876 | Platelet sensitization by LDL | 7,9 |
| TC | M17946 | Valine, leucine and isoleucine biosynthesis | 1,6,9 |
|  | PC:M1+ | (Chaperone) | 3 |
|  | Cerebellum:M4+ | (Response to wounding) | 9 |
|  | Adipose:M1+ |  | 8 |
|  | Omental:M1+ |  | 3 |
|  | rctm1111 | Signal transduction by L1 | 3 |
| TG | rctm1276 | Tight junction interactions | 1,6,8,9 |
|  | rctm0589 | Initial triggering of complement | 1 |
|  | rctm0235 | Cholesterol biosynthesis | 2 |
|  | M18155 | Insulin signaling pathway | 1 |
|  | Blood:M1+ | (Carbohydrate metabolism) | 1,6 |
|  | rctm0225 | Cell-cell junction organization | 1,6,8 |
|  | Blood:M3+ | (Transferase activity, transferring glycosyl groups) | 1 |
|  | M7146 | Classical complement pathway | 1 |
|  | rctm0059 | Activation of Gene Expression by SREBP (SREBF) | 2 |
|  | M917 | Complement pathway | 1 |
|  | M5872 | Steroid biosynthesis | 2 |
|  | Omental:M2+ | (hemopoietic or lymphoid organ development) | 8 |
|  | M2164 | Leukocyte transendothelial migration | 1 |

Note: \*: The method column represents in which methods the MSEA of the pathways is significant with
FDR < 10%. Number 1 to 9 represent: adipose eSNP (1), blood eSNP (2), brain eSNP (3), human aortic
endothelial cells (HAEC) eSNP (4), liver eSNP (5), all eSNP (6), Distance (7), Regulome (8), and Combined (9), respectively. <sup>+</sup>: Co-expression modules. The statistically overrepresented Gene Ontologies satisfying $p < 0.01$ in Fisher's exact test after Benjamini-Hochberg correction within the modules are listed in the parentheses. PC: prefrontal cortex. #: The column tells whether the trait-specific pathways can also be detected as trait-specific ones in either Metabochip and/or iGSEA.

### Replication of lipid-associated pathways using additional dataset and method

To replicate our results from the analysis of GLGC GWAS datasets, we utilized an additional lipid genetic association dataset based on a MetaboChip lipid association study (15) which involved individuals independent of those included in GLGC. The gene sets detected using this independent dataset highly overlapped with those from the GLGC dataset (**Table 1**; **Supplemental Figure S2**; overlapping p values < 10^−20^ by Fisher’s exact test). We also utilized a different pathway analysis method iGSEA (53) and again many of the gene sets were found to be reproducible (**Table 1**; **Supplemental Figure S2**; overlapping p values < 10^−20^).

### Construction of non-overlapping gene supersets for lipid traits

As the knowledge-based pathways and data-driven co-expression modules used in our analysis can converge on similar functional gene units, some of the lipid-associated gene sets have redundancies. We therefore merged overlapping pathways to derive independent, non-overlapping gene sets associated lipid traits. For the 39 shared pathways/co-expression modules across the four lipid traits described earlier, we merged and functionally categorized them into five independent supersets (**Table 1; Table 3**). For the significant gene sets for each lipid trait, we merged them into 17, 16, 18, and 14 supersets for HDL, LDL, TC, and TG, respectively (**Table 3; Supplemental Table S5**), and confirmed that the merged supersets still showed significant association with the corresponding lipid traits in a second round of MSEA (p < 0.05 after Bonferroni correction for the number of supersets tested; **Table 3**).

**Table 3.** Supersets shared by four lipid traits and key driver genes.

| Supersets | No. Genes | Methods<sup>a</sup> |  |  |  | Top Adipose KDs | Top Liver KDs |
| --- | --- | --- | --- | --- | --- | --- | --- |
|  |  | HDL | LDL | TC | TG |  |  |
| Lipid metabolism | 793 | 1,2,3,5 | 1,2,3,5 | 1,2,3,5 | 1,2,3,5 | APOH, ABCB11, F2, ALB, APOA5, APOC4, DMGDH, SERPINC1, APOF, HADHB, ETFDH, KLKB1 | HMGS1, FDFT1, FADS1, DHCR7, ACAT2, ACSS2 |
| Protein catabolism | 253 | 1,3,4,5,6, 7,8,9 | 1,3,5,6 | 1,3,5,6,9 | 1,3,5,6,8 | PSMB9 | PSMB9 |
| Interferon Signaling | 171 | 1,3,5,7,8, 9 | 1,2,3,5,6, 7,8,9 | 1,2,3,5,6, 7,8,9 | 1,2,3,5,6, 8,9 | NUP210 | MX1, ISG15, MX2, IFI44, EPSTI1 |
| Autoimmune/ Immune activation | 152 | 1,3,4,5,6, 7,8,9 | 1,2,3,4,5, 6,7,8,9 | 1,2,3,4,5, 6,7,8,9 | 1,2,3,4,5, 6,7,8,9 | HLA-DMB, HCK, SYK, CD86 | HLA-DMB, CCL5, HLA-DQA1 |
| Visual transduction | 86 | 7,9 | 7,8,9 | 7,8,9 | 7,8,9 | - | - |

**Note:** <sup>a</sup> The method column represents in which methods the MSEA of the pathways is significant with Bonferroni-adjusted $P < 0.05$ . Number 1 to 9 represent: adipose eSNP, blood eSNP, brain eSNP, haec eSNP, liver eSNP, all eSNP, Distance, Regulome, and Combined, respectively.

### Identification of central regulatory genes in the lipid-associated supersets

Subsequently, we performed a key driver analysis (KDA; **Figure 1**) to identify potential regulatory genes or key drivers (KDs) that may regulate genes associated with each lipid trait using Bayesian networks constructed from genetic and gene expression datasets of multiple tissues (detailed in Methods; full KD list in **Supplemental Table S6**). The top adipose and liver KDs for the shared supersets of all four lipid traits and the representative Bayesian subnetworks are shown in **Figure 2**.

**Figure 2.**
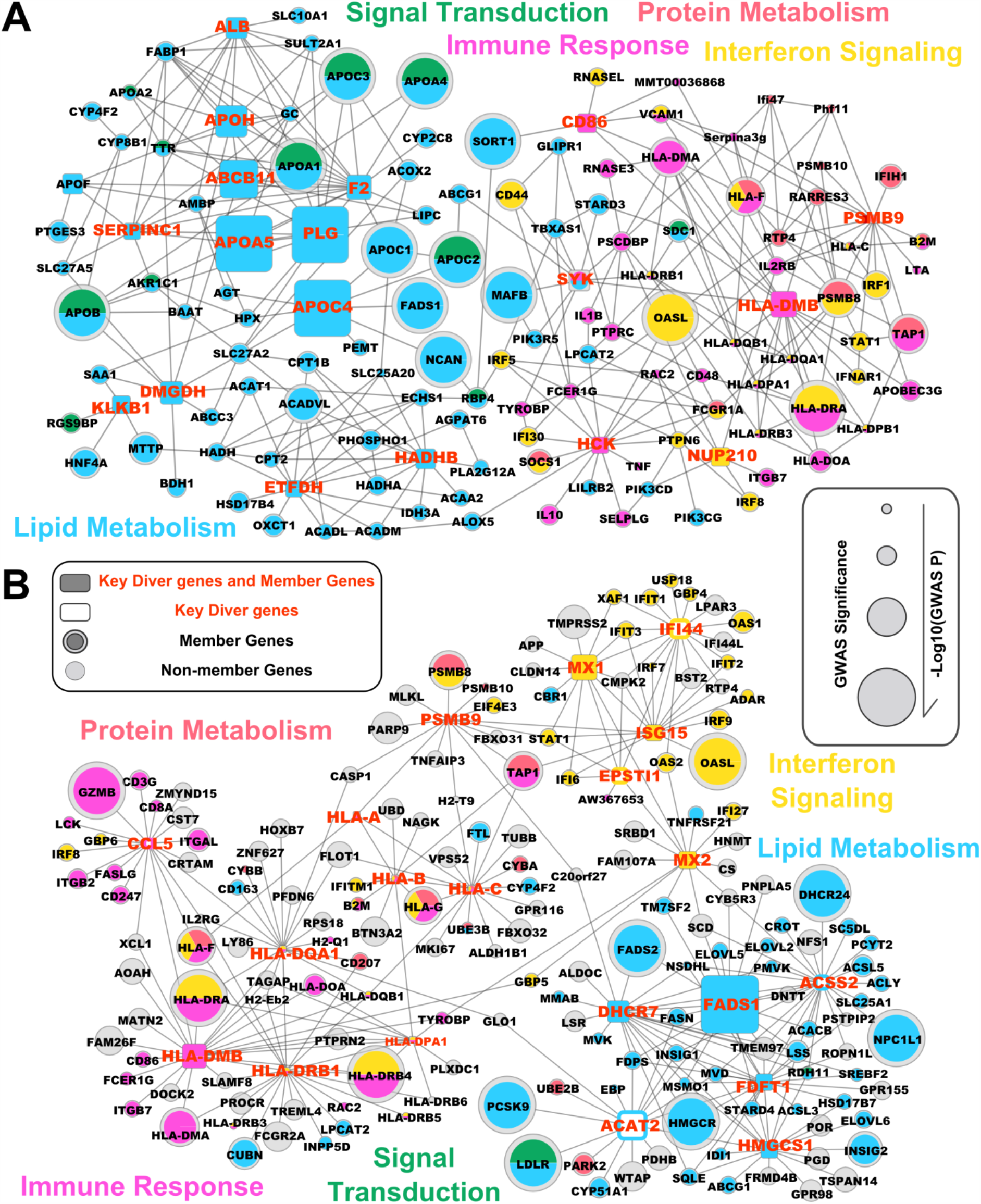
Common KDs and their neighboring genes in the shared lipid-associated subnetworks. A) Adipose KDs and subnetworks. B) Liver KDs and subnetworks. The subnetworks shared by HDL, LDL, TC, and TG are depicted by different colors according to the difference in their functional categories. Nodes are the KDs and their adjacent regulatory partner genes, with KDs depicted as larger nodes. Only network edges that were present in at least two independent network studies were included. The node size corresponds to the GWAS significance.

In adipose tissue (**Figure 2A**), the top KDs for the ‘lipid metabolism’ subnetwork include well-known lipoproteins and ATP-binding cassette (ABC) family members that are responsible for lipid transport, such as *APOF, APOA5* and *ABCB11*. We also found several KDs that are less known to be associated with lipid metabolism, particularly *F2* (coagulation Factor II or thrombin). For the ‘autoimmune/immune activation’ subnetwork, *CD86, HCK*, and *HLA-DMB* were identified as KDs. *PSMB9* was a KD for the ‘protein catabolism’ subnetwork, whereas *NUP210* is central for the ‘interferon signaling’ subnetwork. Moreover, the *SYK* gene is a shared KD between ‘lipid metabolism’ and ‘autoimmune/immune activation’.

In the liver (**Figure 2B**), the top KDs for the ‘lipid metabolism’ subnetwork are enzymes involved in lipid and cholesterol biosynthesis and metabolism, such as *FADS1* (fatty acid desaturase 1), *FDFT1* (farnesyl-diphosphate farnesyltransferase 1), *HMGCS1* (3-hydroxy-3-methylglutaryl-CoA synthase 1), and *DHCR7* (7-dehydrocholesterol reductase). We also identified more KDs for the ‘interferon signaling’ subnetwork in the liver compared to the adipose tissue, with *MX1, MX2, ISG15, IFI44*, and *EPSTI1* being central to the subnetwork. Similar to the adipose network, *PSMB9* and *HLA-DMB* were also identified as KDs for ‘protein catabolism’ and ‘autoimmune/immune activation’ subnetworks in liver, respectively. We did not detect key driver genes for the ‘visual transduction’ subnetwork in either tissue, possibly as a result that the networks of liver and adipose tissues did not capture gene-gene interactions important for this subnetwork.

In addition to the KDs for the subnetworks shared across lipid traits as discussed above, we identified tissue-specific KDs for individual lipid traits (**Supplemental Table S6**). In adipose, *PANK1* and H2B histone family members were specific for the HDL subnetworks (**Figure 3A**); *HIPK2* and *FAU* were top KDs for the LDL subnetworks (**Figure 3B**); genes associated with blood coagulation such as *KNG1* and *FGL1* were KDs for the TC and TG subnetworks (**Figure 3C-3D**). Interestingly, genes related to insulin resistance, *PPARG* and *FASN*, were KDs for both HDL and TG subnetworks. Similarly, trait specific KDs and subnetworks were also detected in the liver; 37 KDs were identified for the TG subnetwork including *ALDH3B1* and *ORM2*, whereas *AHSG, FETUB, ITIH1, HP*, and *SERPINC1* were KDs found in the LDL subnetwork. We note that most of the KDs are themselves not necessarily GWAS hits but are surrounded by significant GWAS genes. For example, gene *F2* is centered by many GWAS hits in the adipose subnetwork (*APOA4, APOC3, APOA5, LIPC*, etc.; **Figure 2; Supplemental Figure S3**). The observation of GWAS hits being peripheral nodes in the network is consistent with previous findings from our group and others (62-67), and again supports that important regulators may not necessarily harbor common variations due to evolutionary constraints.

**Figure 3.**
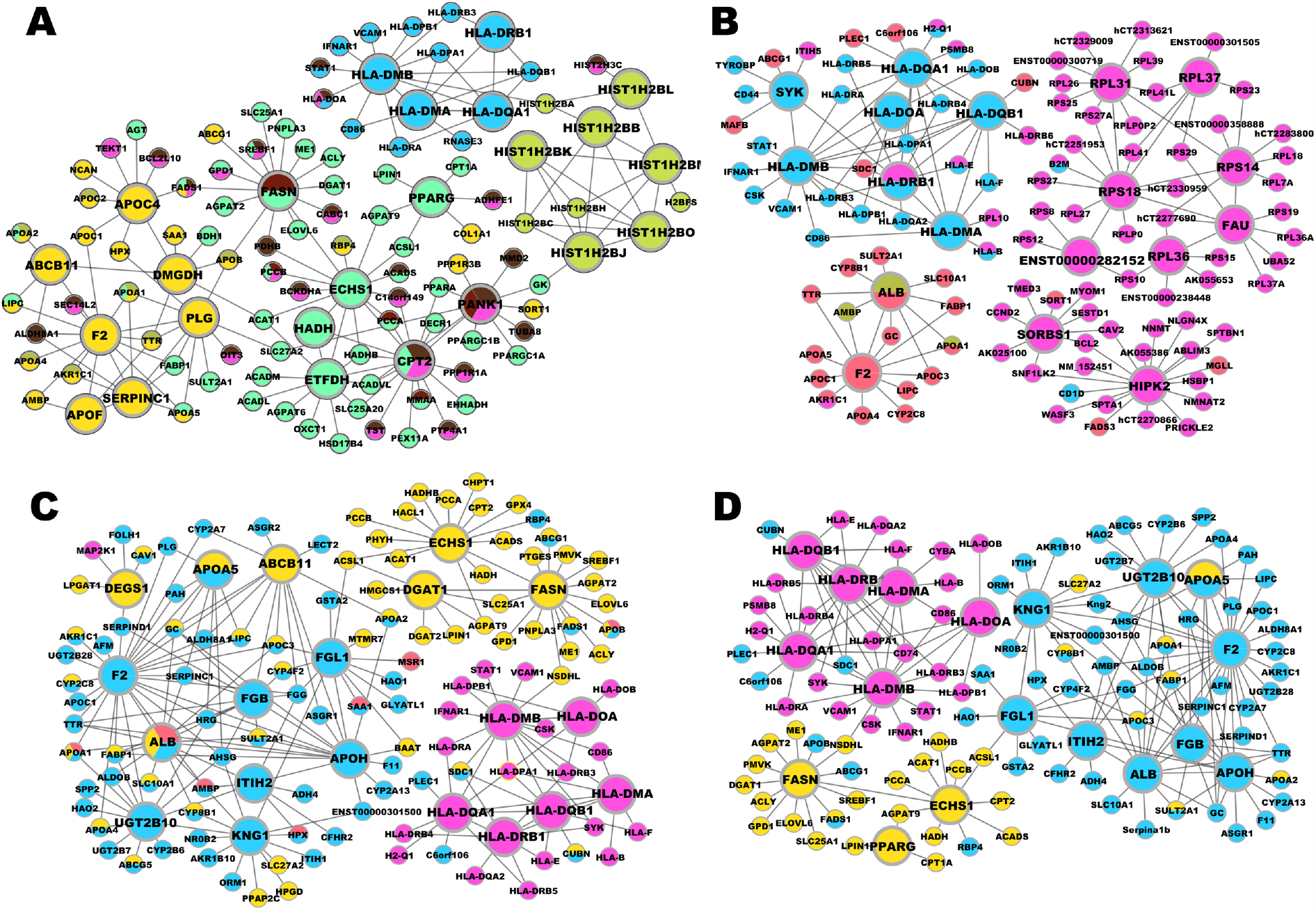
Adipose KDs and subnetworks for each lipid trait. Panel (A)-(D) represent HDL, LDL, TC, and TG subnetworks. Nodes are the KDs and their adjacent regulatory partner genes, with KDs depicted as larger nodes. The yellow color signifies networks associated with interferon signaling, blue with lipid metabolism, pink with immune response, green with protein metabolism, red with lipoprotein metabolism and brown with fatty acid oxidation.

### Experimental validation of *F2* KD subnetworks in 3T3-L1 and C3H10T1/2 adipocytes

Taking into account that the *F2* gene is surrounded by various significant GWAS hits within its subnetwork, we aimed to validate the role of the *F2* gene subnetwork in lipid regulation through siRNA-mediated knockdown experiments in two adipocyte cell lines (3T3-L1 and C3H10T1/2) to ensure reproducibility and robustness of our results. We found that *F2* gene expression was low in preadipocytes for both cells lines, but gradually increased during adipogenesis. In fully differentiated adipocytes between day 8 and day 10, *F2* gene expression level was higher than preadipocytes by 12-fold and 6-fold for 3T3-L1 and C3H10T1/2 lines, respectively (**Figure 4A; 4B)**. When treated with *F2* siRNA, both adipocyte cell lines showed a significant decrease (p < 0.01) in lipid accumulation based on Oil red O staining, as compared with controls treated with scrambled siRNA (**Figure 4C; 4D**). Subsequently, we tested the effect of *F2* gene siRNA knockdown on ten neighbors of the *F2* gene in the adipose network (selected from **Figure 2A**). With 60% knockdown efficiency of *F2* siRNA in the 3T3-L1 adipocytes, seven *F2* network neighbors (*Abcb11, Apoa5, Apof, Fabp1, Lipc, Gc* and *Proc*) exhibited significant changes in expression levels (**Figure 4E**). With 74 % knockdown efficiency of *F2* in C3H10T1/2 adipocytes, six *F2* network neighbors (*Abcb11, Apoa5, Apof, Fabp1, Lipc*, and *Plg*) showed significant changes in expression levels (**Figure 4F**). Several of these genes are involved in lipoprotein transport and fatty acid uptake. In contrast, none of the four negative controls (random genes not in F2 network neighborhood) showed significant changes in their expression levels for 3T3-L1 cell line. However, one negative control gene (*Snrpb2*) did change in the C3H10T1/2 cell line. These results overall support our computational predictions on the structures of *F2* gene subnetworks.

**Figure 4.**
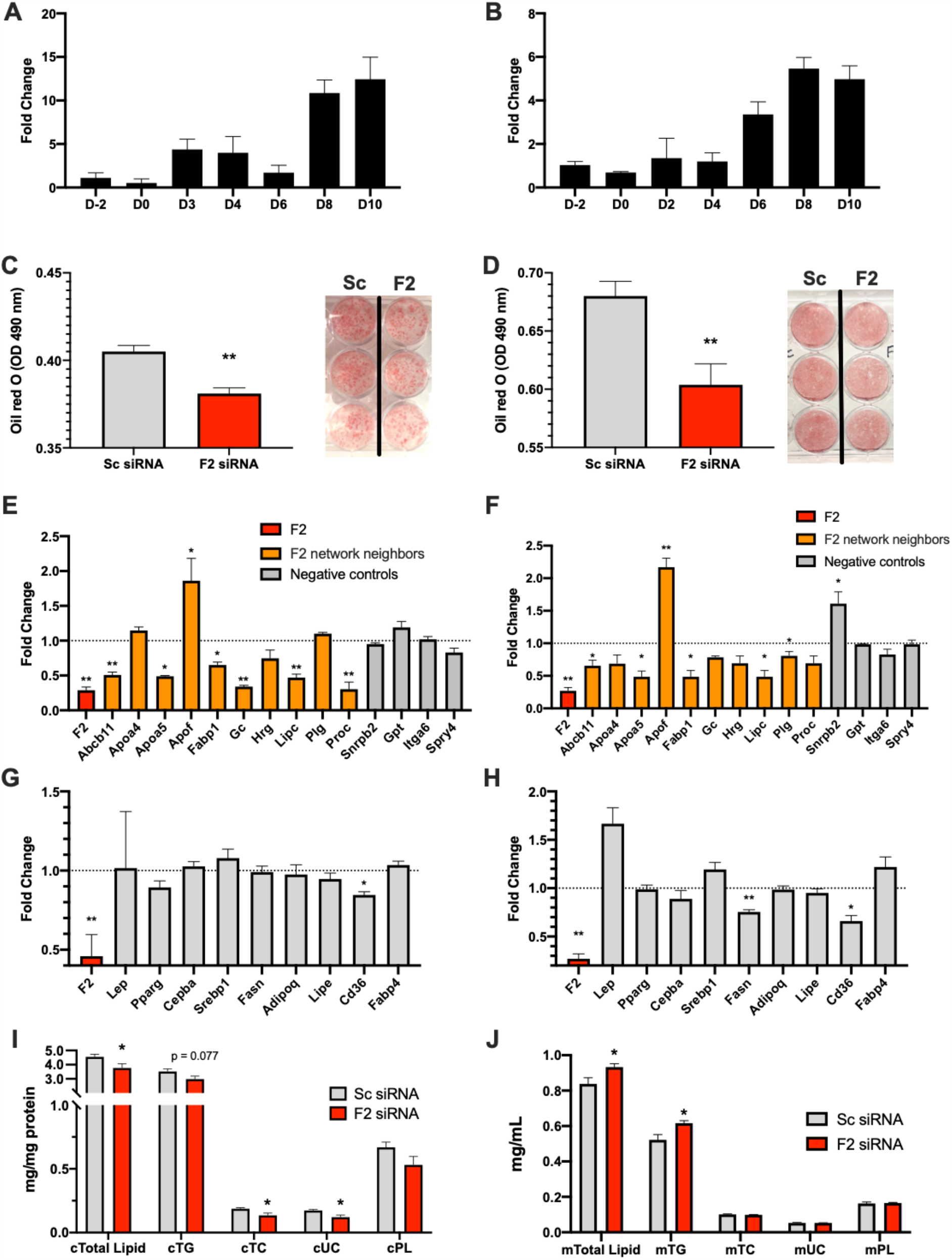
Validation of *F2*’s predicted subnetwork and regulatory role in adipocytes. A, B) Time course of *F2* expression during adipocyte differentiation in 3T3-L1 cells (A) and C3H10T1/2 cells (B). D-2, D0, D2, D3, D4, D6, D8, D10 indicate two days before initiation of differentiation, day 0, day 2, day 3, day 4, day 6, day 8, and day 10 of differentiation, respectively. Sample size n=2-3/time point. C, D) Visualization and quantification (OD value) of lipid accumulation by Oil red O staining in 3T3-L1 adipocytes (C) and C3H10T1/2 adipocytes (D). Sample size n = 5-8/group for adipocytes. E, F) Fold change of expression level for *F2* adipose subnetwork genes and negative control genes after siRNA knockdown. At day 7 of differentiation of 3T3-L1 and day 5 and day 7 of differentiation of C3H10T1/2, adipocytes were transfected with *F2* siRNA for the knockdown experiments. Ten *F2* neighbors were randomly selected from the first and second level neighboring genes of *F2* in adipose network. Four negative controls were randomly selected from the genes not directly connected to *F2* in adipose network. G, H) The fold changes of adipokine/adipogenesis-related genes in 3T3-L1 (G) and in C3H10T1/2 (H). Gene expression levels were determined by RT-qPCR, normalized to Beta actin. The fold changes were relative to scrambled siRNA control. Sample size n=4/group. I, J) Lipid profiles: Total Lipid, Triglyceride (TG), Total Cholesterol (TC), Unesterified Cholesterol (UC) and Phospholipid (PL) in C3H10T1/2 cells (I) and in media (J). Total Lipid was estimated using the sum of the four lipids (TG, TC, UC, PL). Intracellular lipids plotted in (I) were normalized to total cellular protein quantity. Extracellular lipids plotted in (J) are presented as lipid quantity in 1 mL of collected media. Sample size n = 6/group. Results represent mean ± s.e.m. Statistical significance was determined by two-sided Student’s t-test (*p < 0.05 and **p < 0.01).

Next, we measured expression levels of genes related with adipogenesis (*Pparg, Cepba, Srepb1, Fasn*), lipolysis (*Lipe*), fatty acid transport (*Cd36, Fabp4*), and other adipokines following *F2* siRNA treatment. We found no change in the expression of most of the tested genes, with the exception of *Fasn* (in C3H10T1/2), important in the formation of long chain fatty acids, and *Cd36* (in both 3T3-L1 and C3H10T1/2), which encodes fatty acid translocase facilitating fatty acid uptake. *Cd36* expression was decreased by 15 % in 3T3-L1 cells (**Figure 4G**) and 35% in C3H10T1/2 cells (**Figure 4H**) (p < 0.05) and *Fasn* expression was decreased by 25% (**Figure 4H**) (p < 0.01) in C3H10T1/2 cells compared to control. The decreases in *Cd36* and *Fasn* after *F2* knockdown suggest that fatty acid synthesis and uptake by adipocytes are compromised, which could contribute to alterations in circulating lipid levels.

We subsequently measured the lipid contents within the cells and in the media of C3H10T1/2 adipocytes. Following *F2* siRNA treatment, we found significant decreases in total intracellular lipid levels (cTotal Lipid), total cholesterol (cTC), and unesterified cholesterol (cUC), as well as a non-significant trend for decreased triglycerides (cTG) (**Figure 4I**). By contrast, in the culture media, there were significant increases in the total lipid levels (mTotal Lipid) and triglycerides (mTG) following *F2* siRNA treatment (**Figure 4J**). These results support that *F2* knockdown led to decreased intracellular lipids and increased extracellular lipids, agreeing with the overall decreased expression of *F2* network neighbor genes involved in lipid transport and uptake.

### The association between the lipid subnetworks and human diseases

Epidemiological studies consistently show that plasma lipids are closely associated with human complex diseases. For example, high TC and LDL levels are associated with increased risk of cardiovascular disease (CVD). Here, we examined the association between the lipid subnetworks identified in our study and four human complex diseases, namely, Alzheimer’s disease, CVD, T2D, and cancer (Materials and Methods). We found that the gene supersets identified for each lipid traits were significantly enriched for GWAS candidate genes reported by GWAS catalog for the four diseases at Bonferroni-corrected p < 0.05 (**Figure 5; Supplemental Table S7**). The superset ‘lipid metabolism’, which was shared across lipid traits, was associated with Alzheimer’s disease and CVD. When trait-specific subnetworks were considered, those associated with TC, LDL, and TG had more supersets associated with CVD compared to those associated with HDL, a finding consistent with recent reports (15, 68, 69). In addition, supersets of each lipid trait, except HDL, were also found to be significantly associated with cancer, whereas supersets associated with HDL, LDL, and TG but not TC, were linked to T2D.

**Figure 5.**
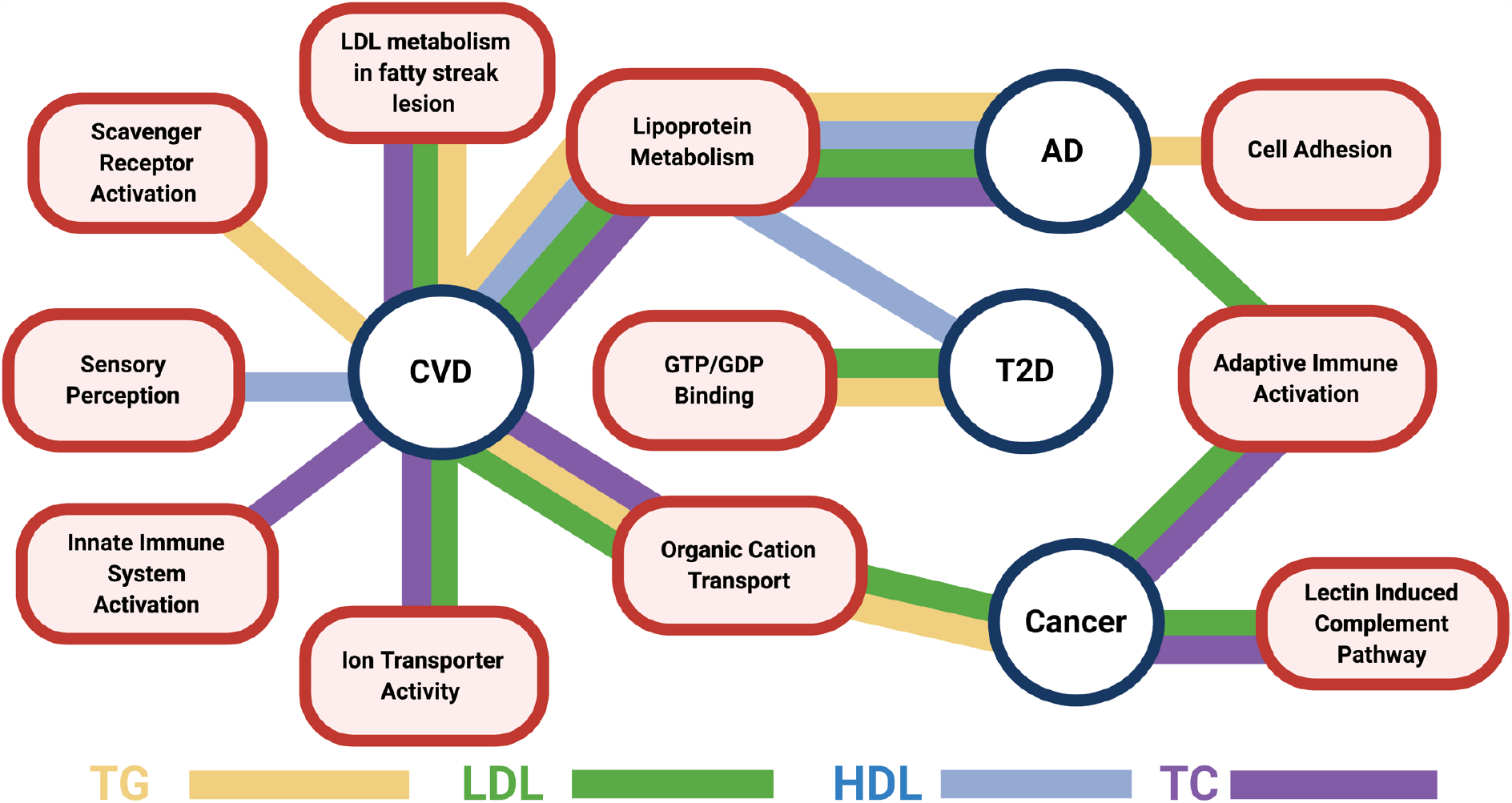
The associations between lipid-associated supersets and human complex diseases. The edges represent the associations between supersets for the specific lipid classes matched by color and diseases (p value < 0.05; Fisher exact test with Bonferroni correction). AD: Alzheimer’s disease; CVD: cardiovascular diseases; T2D: type 2 diabetes.

## Discussion

To gain comprehensive insights into the molecular mechanisms of lipid traits that are important for numerous common complex diseases, we leveraged the large volume of genomic datasets and performed a data-driven multi-omics study combining genetic association signals from large lipid GWAS, tissue-specific eQTLs, ENCODE functional data, known biological pathways, and gene regulatory networks. We identified diverse sets of biological processes, guided by their tissue-specific gene-gene interactions, to be associated with individual lipid traits or shared across lipid traits. Many of the lipid associated gene sets were significantly linked to multiple complex diseases including CVD, T2D, cancer, and Alzheimer’s disease. More importantly, we elucidated tissue-specific gene-gene interactions among the gene sets and identified both well characterized and novel KDs for these lipid-associated processes. We further experimentally validated a novel adipose lipid regulator, *F2*, in two different adipocyte cell lines. Our results offer new insight into the molecular regulation of lipid metabolism, which would not have been possible without the systematic integration of diverse genetic and genomic datasets.

We identified shared pathways associated with all four lipid traits, including ‘lipid metabolism’ and ‘autoimmune/immune activation’, which have been consistently linked to lipid phenotypes, as well as additional pathways such as ‘interferon signaling’, ‘protein catabolism’, and ‘visual transduction’. Interferon factors have previously been linked to lipid storage attenuation and differentiation in human adipocytes (70). Protein catabolism has only recently been identified to be important in regulating lipid metabolism through the PSMD9 protein, which had no previously known function but was shown to cause significant alterations in lipid abundance in both a gain of function and loss of function study in mice (71). The ‘visual transduction’ superset contains retinol-binding proteins, which are carrier proteins involved retinol transport, and play key roles in gene expression regulation and developmental processes (72). ‘visual transduction’ also shares lipoprotein genes with ‘lipid metabolism’, suggesting that retinol-related signal transduction is intimately linked to lipoprotein transport and hence plasma lipid levels.

Furthermore, our results indicate that the trait-specific supersets are tissue-specific. For example, most TG-specific pathways were found to be significant when adipose eSNPs were used, and complement and insulin signaling pathways in the adipose tissue were specific for TG. This is in line with adipose tissue functioning as the major storage site for TG and the regulatory role of immune system and insulin signaling in adipocyte functions and fat storage (73). We also found five HDL-specific pathways, most of which are associated with glucose, lipid, and amino acid metabolism, and were signals derived from liver eSNPs. As HDL acts as the major vehicle for transporting cholesterol to the liver for excretion and catabolism, the critical role of the liver as well as the connections between major metabolic pathways in HDL regulation is recapitulated by our analysis. Interestingly, the TC-specific pathways can be only found when brain eSNPs are used. While the brain accounts for 2% of body weight, it contains 23% of TC in the body (74) and deregulated cholesterol trafficking appears to be involved in the pathogenesis of neurodegenerative diseases, such as Parkinson’s and Alzheimer’s disease (75). These tissue- and trait-specific pathways or processes support the unique features of each lipid species and point to tissue-specific targeting strategies to modulate levels of individual lipid traits and the associated diseases.

In addition to detecting trait- and tissue-specific causal pathways for the lipid traits, our study attempted to delineate the interactions between lipid genes and pathways through gene network analysis. Indeed, the tissue-specific gene networks revealed in our study highlight the regulatory connections between lipid genes and pathways, and thus put individual genes in a broader context. The identification of KDs in a network is essential for uncovering key regulatory components and for identifying drug targets and biomarkers for complex diseases (24, 76). Here, we adopted data-driven Bayesian gene regulatory networks that combine various genomic data (54) to detect the central genes in plasma lipid regulation. The power of this data-driven objective approach has been demonstrated recently (24, 55, 65, 66, 77, 78) and is again supported in this study by the fact that many KDs detected are known regulators for lipids or have served as effective drug targets based on the DrugBank database (79). For instance, for the shared ‘lipid metabolism’ subnetwork, four top KDs (*ACAT2, ACSS2, DHCR7*, and *FADS1*), are targeted by at least one FDA approved anti-cholesteremic drug. Another KD, *HMGCS1*, is a rate-limiting enzyme of cholesterol synthesis, and is considered a promising drug target in lipid-associated metabolic disorders (80). These lines of evidence lead us to speculate the other less-studied KDs are also important for lipid regulation.

Among the top network KDs predicted, several including *F2, KLKB1* and *ANXA4* are involved in blood coagulation. A previous study revealed polymorphisms in the anticoagulation genes modify the efficacy of statins in reducing risk of cardiovascular events (81), which in itself is not surprising. However, the intimate relationship between a coagulation gene *F2* and lipid regulation predicted by our analysis is intriguing (**Figure 4)**. We found that the partner genes in the adipose *F2* subnetwork tend to be differentially expressed after *F2* knockdown in both 3T3-L1 and C3H10T1/2 adipocytes, with several of the altered genes (*Apoa5, Apof, Abcb11, Fabp1, Fasn* and *Cd36*) closely associated with cholesterol and fatty acid transport and uptake. We further observed that *F2* knockdown affects lipid storage in adipocytes, with intracellular lipid content decreasing and extracellular lipid content in the media increasing. Interestingly, *F2* expression level is low in preadipocytes and only increases during the late phase of adipocyte differentiation. Our findings support a largely untapped role of *F2* in lipid transport and storage in adipocytes and provide a novel target in the *F2* gene.

In addition to the shared KDs such as *F2* for different lipids, it may be also of value to focus on the trait-specific KDs as numerous studies have revealed these lipid phenotypes play different roles in many human diseases. For example, LDL and TC are important risk factors for CVD (82) and TG has been linked to T2D (83), while the role of HDL in CVD has been controversial (84). We detected 37 genes as TG-specific KDs in liver regulatory subnetworks. Among these, *CP* (ceruloplasmin) and *ALDH3B1* (aldehyde dehydrogenase 3 family, member B1) were clinically confirmed to be associated with T2D (85, 86) while most of the other genes such as *DHODH* and *ANXA4* were less known to be associated with TG and thus may serve as novel targets. In adipose tissue, genes important for insulin resistance and diabetes such as *PPARG* and *FASN* were found to be KDs for TG, further supporting the connection between TG and diabetes. Additionally, *FASN* has been implicated as a KD in numerous studies for non-alcoholic fatty liver disease (67, 78, 87), again highlighting the importance of this gene in common metabolic disorders.

We acknowledge some potential limitations to our study. First, the GWAS datasets utilized are not the most recently conducted and therefore provides the possibility of not capturing the full array of unknown biology. However, despite this our results are consistent with the biology found more recently including overlapping signals in pathways for chylomicron-mediated lipid transport and lipoprotein metabolism (88) as well as more novel findings such as visual transduction pathways. In addition, one of our key drivers *KLKB1*, which was not found to be a GWAS hit in the dataset we utilized, has since been found to pass the genome wide significance threshold in more recent larger GWAS and is a hit on apolipoprotein A-IV concentrations, which is a major component of HDL and chylomicron particles important in reverse cholesterol transport (89). This further exemplifies the robustness of our integrative network approach to find key genes important to disease pathogenesis even when smaller GWAS were utilized.

In summary, we used an integrative genomics framework to leverage a multitude of genetic and genomic datasets from human studies to unravel the underlying regulatory processes involved in lipid phenotypes. We not only detected shared processes and gene regulatory networks among different lipid traits, but also provide comprehensive insight into trait-specific pathways and networks. The results suggest there are both shared and distinct mechanisms underlying very closely related lipid phenotypes. The tissue-specific networks and KDs identified in our study shed light on molecular mechanisms involved in lipid homeostasis. If validated in additional population genetic and mechanistic studies, these molecular processes and genes can be used as novel targets for the treatment of lipid-associated disorders such as CVD, T2D, Alzheimer’s disease and cancers.

## Data Availability

All genomic data utilized in the analysis were previously published and were downloaded from public data repositories. All experimental data were presented in the current manuscript. Mergeomics code is available at R Bioconductor DOI: <u>10.18129/B9.bioc.Mergeomics</u>.

## Acknowledgements

We would like to thank Dr. Aldons J. Lusis in the Department of Human Genetics, UCLA for valuable discussions during the preparation of the manuscript. We would also like to thank Gajalakshmi Ramanathan for technical support with the in vitro validation analysis and Dr. Marcus Tol and Dr. Peter Tontonoz in the Department of Pathology and Laboratory Medicine in the David Geffen School of Medicine at UCLA for providing the C3H10T1/2 adipocyte cell lines. XY is supported by the National Institutes of Health Grants R01 DK104363 and R01 DK117850.

## Author contributions

XY and YZ designed and directed the study. MB, YZ, ISA, ZS, and HL conducted the analyses. VPM contributed analytical methods and tools. MB, ZS, ISA, YZ and XY wrote the manuscript. ISA and IC conducted the validation experiments. All authors edited and approved the final manuscript.

## References

1. Austin MA. Plasma Triglyceride and Coronary Heart-Disease. Arterioscler Thromb. 1991;11(1):2–14.

2. Reitz C, Tang MX, Luchsinger J, Mayeux R. Relation of plasma lipids to Alzheimer disease and vascular dementia. Arch Neurol-Chicago. 2004;61(5):705–14.

3. Di Paolo G, Kim TW. Linking lipids to Alzheimer’s disease: cholesterol and beyond. Nat Rev Neurosci. 2011;12(5):284–96.

4. Muoio DM, Newgard CB. Molecular and metabolic mechanisms of insulin resistance and β-cell failure in type 2 diabetes. Nature reviews Molecular cell biology. 2008;9(3):193.

5. Zhang F, Du G. Dysregulated lipid metabolism in cancer. World journal of biological chemistry. 2012;3(8):167.

6. Zhang BB, Zhou GC, Li C. AMPK: An Emerging Drug Target for Diabetes and the Metabolic Syndrome. Cell Metab. 2009;9(5):407–16.

7. Libby P, Ridker PM, Hansson GK. Progress and challenges in translating the biology of atherosclerosis. Nature. 2011;473(7347):317–25.

8. Tabas I, Glass CK. Anti-Inflammatory Therapy in Chronic Disease: Challenges and Opportunities. Science. 2013;339(6116):166–72.

9. Heller DA, Defaire U, Pedersen NL, Dahlen G, Mcclearn GE. Genetic and Environmental-Influences on Serum-Lipid Levels in Twins. New Engl J Med. 1993;328(16):1150–6.

10. Kathiresan S, Melander O, Guiducci C, Surti A, Burtt NP, Rieder MJ, et al. Six new loci associated with blood low-density lipoprotein cholesterol, high-density lipoprotein cholesterol or triglycerides in humans. Nat Genet. 2008;40(2):189–97.

11. van Dongen J, Willemsen G, Chen WM, de Geus EJ, Boomsma DI. Heritability of metabolic syndrome traits in a large population-based sample. J Lipid Res. 2013;54(10):2914–23.

12. Teslovich TM, Musunuru K, Smith AV, Edmondson AC, Stylianou IM, Koseki M, et al. Biological, clinical and population relevance of 95 loci for blood lipids. Nature. 2010;466(7307):707–13.

13. Kathiresan S, Manning AK, Demissie S, D’Agostino RB, Surti A, Guiducci C, et al. A genome-wide association study for blood lipid phenotypes in the Framingham Heart Study. Bmc Med Genet. 2007;8.

14. Willer CJ, Sanna S, Jackson AU, Scuteri A, Bonnycastle LL, Clarke R, et al. Newly identified loci that influence lipid concentrations and risk of coronary artery disease. Nat Genet. 2008;40(2):161–9.

15. Willer CJ, Schmidt EM, Sengupta S, Peloso GM, Gustafsson S, Kanoni S, et al. Discovery and refinement of loci associated with lipid levels. Nat Genet. 2013;45(11):1274–83.

16. Klarin D, Damrauer SM, Cho K, Sun YV, Teslovich TM, Honerlaw J, et al. Genetics of blood lipids among∼ 300,000 multi-ethnic participants of the Million Veteran Program. Nature genetics. 2018;50(11):1514.

17. Hoffmann TJ, Theusch E, Haldar T, Ranatunga DK, Jorgenson E, Medina MW, et al. A large electronic-health-record-based genome-wide study of serum lipids. Nature genetics. 2018;50(3):401.

18. Zhong H, Yang X, Kaplan LM, Molony C, Schadt EE. Integrating Pathway Analysis and Genetics of Gene Expression for Genome-wide Association Studies. Am J Hum Genet. 2010;86(4):581–91.

19. Lonsdale J, Thomas J, Salvatore M, Phillips R, Lo E, Shad S, et al. The genotype-tissue expression (GTEx) project. Nature genetics. 2013;45(6):580.

20. Boyle AP, Hong EL, Hariharan M, Cheng Y, Schaub MA, Kasowski M, et al. Annotation of functional variation in personal genomes using RegulomeDB. Genome Res. 2012;22(9):1790–7.

21. MacNeil LT, Walhout AJM. Gene regulatory networks and the role of robustness and stochasticity in the control of gene expression. Genome Res. 2011;21(5):645–57.

22. Wang K, Li MY, Bucan M. Pathway-based approaches for analysis of genomewide association studies. Am J Hum Genet. 2007;81(6):1278–83.

23. Zhong H, Beaulaurier J, Lum PY, Molony C, Yang X, MacNeil DJ, et al. Liver and Adipose Expression Associated SNPs Are Enriched for Association to Type 2 Diabetes. Plos Genet. 2010;6(5).

24. Makinen VP, Civelek M, Meng Q, Zhang B, Zhu J, Levian C, et al. Integrative genomics reveals novel molecular pathways and gene networks for coronary artery disease. Plos Genet. 2014;10(7):e1004502.

25. Baranzini SE, Galwey NW, Wang J, Khankhanian P, Lindberg R, Pelletier D, et al. Pathway and network-based analysis of genome-wide association studies in multiple sclerosis. Hum Mol Genet. 2009;18(11):2078–90.

26. Jia PL, Wang LL, Meltzer HY, Zhao ZM. Common variants conferring risk of schizophrenia: A pathway analysis of GWAS data. Schizophr Res. 2010;122(1-3):38–42.

27. Willer CJ, Schmidt EM, Sengupta S, Peloso GM, Gustafsson S, Kanoni S, et al. Discovery and refinement of loci associated with lipid levels. Nat Genet. 2013;45(11):1274-+.

28. Joshi-Tope G, Gillespie M, Vastrik I, D’Eustachio P, Schmidt E, de Bono B, et al. Reactome: a knowledgebase of biological pathways. Nucleic Acids Res. 2005;33:D428–D32.

29. Ogata H, Goto S, Sato K, Fujibuchi W, Bono H, Kanehisa M. KEGG: Kyoto Encyclopedia of Genes and Genomes. Nucleic Acids Res. 1999;27(1):29–34.

30. TA HLSPJHREMJCFM. Potential etiologic and functional implications of genome-wide association loci for human diseases and traits. Proc Natl Acad Sci U S A. 2009;106(23):9362–7.

31. Emilsson V, Thorleifsson G, Zhang B, Leonardson AS, Zink F, Zhu J, et al. Genetics of gene expression and its effect on disease. Nature. 2008;452(7186):423–U2.

32. Derry JMJ, Zhong H, Molony C, MacNeil D, Guhathakurta D, Zhang B, et al. Identification of Genes and Networks Driving Cardiovascular and Metabolic Phenotypes in a Mouse F2 Intercross. Plos One. 2010;5(12).

33. Schadt EE, Molony C, Chudin E, Hao K, Yang X, Lum PY, et al. Mapping the genetic architecture of gene expression in human liver. Plos Biol. 2008;6(5):1020–32.

34. Greenawalt DM, Dobrin R, Chudin E, Hatoum IJ, Suver C, Beaulaurier J, et al. A survey of the genetics of stomach, liver, and adipose gene expression from a morbidly obese cohort. Genome Res. 2011;21(7):1008–16.

35. Fehrmann RSN, Jansen RC, Veldink JH, Westra HJ, Arends D, Bonder MJ, et al. Trans-eQTLs Reveal That Independent Genetic Variants Associated with a Complex Phenotype Converge on Intermediate Genes, with a Major Role for the HLA. PLoS genetics. 2011;7(8).

36. Wang SS, Schadt EE, Wang H, Wang XP, Ingram-Drake L, Shi W, et al. Identification of pathways for atherosclerosis in mice - Integration of quantitative trait locus analysis and global gene expression data. Circ Res. 2007;101(3):E11–E30.

37. Yang X, Schadt EE, Wang S, Wang H, Arnold AP, Ingram-Drake L, et al. Tissue-specific expression and regulation of sexually dimorphic genes in mice. Genome Res. 2006;16(8):995–1004.

38. Tu ZD, Keller MP, Zhang CS, Rabaglia ME, Greenawalt DM, Yang X, et al. Integrative Analysis of a Cross-Loci Regulation Network Identifies App as a Gene Regulating Insulin Secretion from Pancreatic Islets. PLoS genetics. 2012;8(12).

39. Nica AC, Parts L, Glass D, Nisbet J, Barrett A, Sekowska M, et al. The Architecture of Gene Regulatory Variation across Multiple Human Tissues: The MuTHER Study. PLoS genetics. 2011;7(2).

40. Romanoski CE, Che N, Yin F, Mai N, Pouldar D, Civelek M, et al. Network for Activation of Human Endothelial Cells by Oxidized Phospholipids. Circ Res. 2011;109(5):E27–U52.

41. Romanoski CE, Che N, Yin F, Mai N, Pouldar D, Civelek M, et al. Network for activation of human endothelial cells by oxidized phospholipids: a critical role of heme oxygenase 1. Circ Res. 2011;109(5):e27–41.

42. Dixon AL, Liang L, Moffatt MF, Chen W, Heath S, Wong KC, et al. A genome-wide association study of global gene expression. Nature genetics. 2007;39(10):1202–7.

43. Fehrmann RS, Jansen RC, Veldink JH, Westra HJ, Arends D, Bonder MJ, et al. Trans-eQTLs reveal that independent genetic variants associated with a complex phenotype converge on intermediate genes, with a major role for the HLA. Plos Genet. 2011;7(8):e1002197.

44. Nica AC, Parts L, Glass D, Nisbet J, Barrett A, Sekowska M, et al. The architecture of gene regulatory variation across multiple human tissues: the MuTHER study. Plos Genet. 2011;7(2):e1002003.

45. Montgomery SB, Sammeth M, Gutierrez-Arcelus M, Lach RP, Ingle C, Nisbett J, et al. Transcriptome genetics using second generation sequencing in a Caucasian population. Nature. 2010;464(7289):773–U151.

46. Stranger BE, Montgomery SB, Dimas AS, Parts L, Stegle O, Ingle CE, et al. Patterns of Cis Regulatory Variation in Diverse Human Populations. PLoS genetics. 2012;8(4):272–84.

47. Stranger BE, Nica AC, Forrest MS, Dimas A, Bird CP, Beazley C, et al. Population genomics of human gene expression. Nature genetics. 2007;39(10):1217–24.

48. Dimas AS, Deutsch S, Stranger BE, Montgomery SB, Borel C, Attar-Cohen H, et al. Common Regulatory Variation Impacts Gene Expression in a Cell Type-Dependent Manner. Science. 2009;325(5945):1246–50.

49. Duan S, Huang RS, Zhang W, Bleibel WK, Roe CA, Clark TA, et al. Genetic architecture of transcript-level variation in humans. Am J Hum Genet. 2008;82(5):1101–13.

50. Maher B. ENCODE: The human encyclopaedia. Nature. 2012;489(7414):46–8.

51. Shu L, Zhao Y, Kurt Z, Byars SG, Tukiainen T, Kettunen J, et al. Mergeomics: integration of diverse genomics resources to identify pathogenic perturbations to biological systems. bioRxiv. 2016:036012.

52. Benjamini Y, Hochberg Y. Controlling the False Discovery Rate - a Practical and Powerful Approach to Multiple Testing. J Roy Stat Soc B Met. 1995;57(1):289–300.

53. Zhang KL, Cui SJ, Chang SH, Zhang LY, Wang J. i-GSEA4GWAS: a web server for identification of pathways/gene sets associated with traits by applying an improved gene set enrichment analysis to genome-wide association study. Nucleic Acids Res. 2010;38:W90–W5.

54. Zhu J, Zhang B, Smith EN, Drees B, Brem RB, Kruglyak L, et al. Integrating large-scale functional genomic data to dissect the complexity of yeast regulatory networks. Nature genetics. 2008;40(7):854–61.

55. Wang IM, Zhang B, Yang X, Zhu J, Stepaniants S, Zhang CS, et al. Systems analysis of eleven rodent disease models reveals an inflammatome signature and key drivers. Mol Syst Biol. 2012;8.

56. Yang X, Zhang B, Molony C, Chudin E, Hao K, Zhu J, et al. Systematic genetic and genomic analysis of cytochrome P450 enzyme activities in human liver. Genome Res. 2010;20(8):1020–36.

57. Ye J, Coulouris G, Zaretskaya I, Cutcutache I, Rozen S, Madden TL. Primer-BLAST: a tool to design target-specific primers for polymerase chain reaction. BMC bioinformatics. 2012;13(1):134.

58. Livak KJ, Schmittgen TD. Analysis of relative gene expression data using real-time quantitative PCR and the 2(T)(-Delta Delta C) method. Methods. 2001;25(4):402–8.

59. Folch J, Lees M, Stanley GS. A simple method for the isolation and purification of total lipides from animal tissues. Journal of biological chemistry. 1957;226(1):497–509.

60. Diamante G, Cely I, Zamora Z, Ding J, Blencowe M, Lang J, et al. Systems toxicogenomics of prenatal low-dose BPA exposure on liver metabolic pathways, gut microbiota, and metabolic health in mice. Environment International. 2021;146:06260.

61. Welter D, MacArthur J, Morales J, Burdett T, Hall P, Junkins H, et al. The NHGRI GWAS Catalog, a curated resource of SNP-trait associations. Nucleic acids research. 2013;42(D1):D1001–D6.

62. Goh KI, Cusick ME, Valle D, Childs B, Vidal M, Barabasi AL. The human disease network. P Natl Acad Sci USA. 2007;104(21):8685–90.

63. Makinen VP, Civelek M, Meng QY, Zhang B, Zhu J, Levian C, et al. Integrative Genomics Reveals Novel Molecular Pathways and Gene Networks for Coronary Artery Disease. Plos Genet. 2014;10(7).

64. Shu L, Chan KHK, Zhang G, Huan T, Kurt Z, Zhao Y, et al. Shared genetic regulatory networks for cardiovascular disease and type 2 diabetes in multiple populations of diverse ethnicities in the United States. PLoS genetics. 2017;13(9):e1007040.

65. Zhao Y, Blencowe M, Shi X, Shu L, Levian C, Ahn IS, et al. Integrative Genomics Analysis Unravels Tissue-Specific Pathways, Networks, and Key Regulators of Blood Pressure Regulation. Frontiers in cardiovascular medicine. 2019;6:21.

66. Zhao Y, Jhamb D, Shu L, Arneson D, Rajpal DK, Yang X. Multi-omics integration reveals molecular networks and regulators of psoriasis. BMC systems biology. 2019;13(1):8.

67. Krishnan KC, Kurt Z, Barrere-Cain R, Sabir S, Das A, Floyd R, et al. Integration of multi-omics data from mouse diversity panel highlights mitochondrial dysfunction in non-alcoholic fatty liver disease. Cell systems. 2018;6(1):103-15. e7.

68. Hewing B, Landmesser U. LDL, HDL, VLDL, and CVD prevention: lessons from genetics? Current cardiology reports. 2015;17(7):56.

69. Santos-Gallego CG. HDL: quality or quantity? Atherosclerosis. 2015;243(1):121–3.

70. McGillicuddy FC, Chiquoine EH, Hinkle CC, Kim RJ, Shah R, Roche HM, et al. Interferon γ attenuates insulin signaling, lipid storage, and differentiation in human adipocytes via activation of the JAK/STAT pathway. Journal of Biological Chemistry. 2009;284(46):31936–44.

71. Parker BL, Calkin AC, Seldin MM, Keating MF, Tarling EJ, Yang P, et al. An integrative systems genetic analysis of mammalian lipid metabolism. Nature. 2019;567(7747):187.

72. Zizola C, Frey SK, Jitngarmkusol S, Kadereit B, Yan N, Vogel S. Cellular retinol-binding protein type I (CRBP-I) regulates adipogenesis. Molecular and cellular biology. 2010;30(14):3412–20.

73. Schäffler A, Schölmerich J. Innate immunity and adipose tissue biology. Trends in immunology. 2010;31(6):228–35.

74. Vance JE. MAM (mitochondria-associated membranes) in mammalian cells: lipids and beyond. Biochimica et Biophysica Acta (BBA)-Molecular and Cell Biology of Lipids. 2014;1841(4):595–609.

75. Liu JP, Tang Y, Zhou SF, Toh BH, McLean C, Li H. Cholesterol involvement in the pathogenesis of neurodegenerative diseases. Mol Cell Neurosci. 2010;43(1):33–42.

76. Jeong H, Mason SP, Barabasi AL, Oltvai ZN. Lethality and centrality in protein networks. Nature. 2001;411(6833):41–2.

77. Zhang B, Gaiteri C, Bodea LG, Wang Z, McElwee J, Podtelezhnikov AA, et al. Integrated systems approach identifies genetic nodes and networks in late-onset Alzheimer’s disease. Cell. 2013;153(3):707–20.

78. Kurt Z, Barrere-Cain R, LaGuardia J, Mehrabian M, Pan C, Hui ST, et al. Tissue-specific pathways and networks underlying sexual dimorphism in non-alcoholic fatty liver disease. Biology of sex differences. 2018;9(1):46.

79. Knox C, Law V, Jewison T, Liu P, Ly S, Frolkis A, et al. DrugBank 3.0: a comprehensive resource for ‘Omics’ research on drugs. Nucleic Acids Res. 2011;39:D1035–D41.

80. Yue WW, Oppermann U. High-throughput structural biology of metabolic enzymes and its impact on human diseases. J Inherit Metab Dis. 2011;34(3):575–81.

81. Maitland-van der Zee Ah, Peters BJM, Lynch AI, Boerwinkle E, Arnett DK, Cheng S, et al. The effect of nine common polymorphisms in coagulation factor genes (F2, F5, F7, F12 and F13) on the effectiveness of statins: the GenHAT study. Pharmacogenet Genom. 2009;19(5):338–44.

82. Ference BA, Yoo W, Alesh I, Mahajan N, Mirowska KK, Mewada A, et al. Effect of Long-Term Exposure to Lower Low-Density Lipoprotein Cholesterol Beginning Early in Life on the Risk of Coronary Heart Disease A Mendelian Randomization Analysis. J Am Coll Cardiol. 2012;60(25):2631–9.

83. Guilherme A, Virbasius JV, Puri V, Czech MP. Adipocyte dysfunctions linking obesity to insulin resistance and type 2 diabetes. Nature reviews Molecular cell biology. 2008;9(5):367–77.

84. Voight BF, Peloso GM, Orho-Melander M, Frikke-Schmidt R, Barbalic M, Jensen MK, et al. Plasma HDL cholesterol and risk of myocardial infarction: a mendelian randomisation study. Lancet. 2012;380(9841):572–80.

85. Memisogullari R, Bakan E. Levels of ceruloplasmin, transferrin, and lipid peroxidation in the serum of patients with Type 2 diabetes mellitus. J Diabetes Complicat. 2004;18(4):193–7.

86. Volkmar M, Dedeurwaerder S, Cunha DA, Ndlovu MN, Defrance M, Deplus R, et al. DNA methylation profiling identifies epigenetic dysregulation in pancreatic islets from type 2 diabetic patients. Embo J. 2012;31(6):1405–26.

87. Blencowe M, Karunanayake T, Wier J, Hsu N, Yang X. Network Modeling Approaches and Applications to Unravelling Non-Alcoholic Fatty Liver Disease. Genes. 2019;10(12):966.

88. Chen L, Yao Y, Jin C, Wu S, Liu Q, Li J, et al. Integrative genomic analysis identified common regulatory networks underlying the correlation between coronary artery disease and plasma lipid levels. BMC Cardiovascular Disorders. 2019;19(1):1–10.

89. Lamina C, Friedel S, Coassin S, Rueedi R, Yousri NA, Seppälä I, et al. A genome-wide association meta-analysis on apolipoprotein A-IV concentrations. Human molecular genetics. 2016;25(16):3635–46.

