## Supplemental Materials for "Gene Networks and Pathways for Plasma Lipid Traits via Multi-tissue Multi-omics Systems Analysis"

for

### Supplemental Methods

The overall statistical framework can be divided into three parts, including Marker set enrichment analysis (MSEA), construction of non-overlapping supersets, and key driver analysis (KDA).

### Marker set enrichment analysis (MSEA)

We applied a modified MSEA method [1] to identify biological pathways or co-expression modules associated with lipid traits. In the analysis, we aimed for a test statistic that exhibits a symmetric null distribution to make it easier to approximate results from permutation analysis and that is based on quantiles of lipid association signals to make the test non-parametric. Quantiles in this context refer to the inverted association P-values from a GWAS study. For instance, a quantile point of 95% would mean that the top 5% of association signals are assigned as positive and the rest as negative associations. Importantly, we use multiple quantile cutoffs (70%, 80%, 90%, 99%, 99.9%, and 99.99%) to reduce the need to optimize a specific cutoff and to achieve high overall sensitivity. We adopted a modified $X^2$-test (Chi-square-like) to calculate the enrichment of lipid genetic signals for each of the selected gene sets. The formula is as follows:

$$\chi = \sum_{i=1}^{n} \frac{O_i - E_i}{\sqrt{E_i} + \kappa}$$

where $n$ denotes the number of quantile cutoffs, $O$ and $E$ the observed and expected counts of positive findings, and $\kappa = 1$ denotes the stability parameter to reduce artifacts from low expected counts. The null hypothesis for the enrichment of GWAS signals within a pathway is defined as 'Given the set of all distinct loci from a set of N genes, these loci contain an equal proportion of positive GWAS findings when compared to all the distinct loci from a set of N random genes'. The expected distribution of the test statistic under the null hypothesis can be estimated empirically by randomly shuffling the genes. The approach is robust against LD and other artifacts. The exact shape of the null distribution is dependent on the size of the gene set, and on the mapping between the genes and the loci. To make the enrichment statistics of the gene sets comparable with one another, we adapted a logarithmic method that was originally developed for the self-organizing map. Let $X_0$ denote the vector of simulated test statistics from the permutation analysis for a single gene set, then the transformation algorithm can be expressed as

1) $\alpha = \min(X_0)$, $X_1 = X_0 - \alpha$

2) $\beta = \mathrm{median}(X_1)$, $X_2 = X_1 / \beta$

3) $X_3 = \log(\gamma X_2 + 1)$

4) $\mu = \mathrm{mean}(X_3)$, $\sigma = \mathrm{sd}(X_3)$

5) Evaluate how well $X_3$ approximates $N(\mu, \sigma)$

6) If necessary, try a different $\gamma$ and go back to Step 3.

The parameters from Steps 1-4 can be saved and reapplied to new data, which makes it possible to determine the transformation exclusively based on simulated statistics, and then apply it to the observed test statistic to yield the parametric enrichment score

$$Z_N = \frac{\log(\gamma(\chi - \alpha)\beta^{-1} + 1) - \mu}{\sigma}$$

The rationale for Gaussian approximation is based on the attractive analytical properties of Gaussian distributions. Nevertheless, if the approximation is inaccurate, the results can be biased and lead to erroneous conclusion. In particular, any dependencies between genetic loci tend to elongate the tails of the "true" distribution. For this reason, we also report the frequency of false positive findings from the permutation analysis for each gene set, convert the frequencies into standardized scores $Z_F$ via the inverse cumulative Gaussian density function, and use the mean of the parametric score $Z_N$ and frequency-based scores as the final enrichment measure z:

$$Z_F = N^{-1}(p_F)$$

$$z = (Z_N + Z_F)/2$$

The reported gene set P-value is estimated from the z-score according to the cumulative Gaussian density. False discovery rates were estimated with the method by Benjamini and Hochberg.

**Construction of non-overlapping supersets**

We expected to find redundant gene sets among the top hits as a consequence of the nested and overlapping definitions from multiple pathway databases and co-expression studies. To make the results more meaningful, we constructed non-overlapping supersets that captured the core genes from groups of redundant pathways. For two gene sets A and B with different numbers of member genes, two overlap ratios were calculated: the proportion of genes in A that were also in B ($r_{AB}$), and the proportion of genes in B that were also in A ($r_{BA}$). We chose the formula $r = (r_{AB} \times r_{BA})^{0.5}$ to describe the degree of overlap. Importantly, $r$ is small whenever the sizes of A and B are substantially different, which discourages the merging of nested gene sets. As an additional overlap criterion, we required that Fisher's exact test for the number of shared genes was statistically significant ($P < 0.05$ after Bonferroni correction).

We employed hierarchical clustering to define blocks of overlapping gene sets. First, the overlap matrix of the lipid-associated gene sets was estimated, and all non-significant elements were set to zero. The overlaps were then converted to distances $d = (1 - r)$. Clusters of overlapping gene sets were identified by the hclust() function in the R programming environment with a static cutoff at zero overlap. In the last step, the gene sets within clusters were merged and trimmed into supersets.

A size limit of 200 genes was chosen to trim the raw supersets down to the core genes that were shared across overlapping gene sets. This choice of optimal size was motivated by earlier MSEA analyses [1]. The least shared genes were successively removed until the next removal would have reduced the superset size below the 200-gene limit. Overlap ratios were re-calculated between the trimmed supersets before the next round of hierarchical clustering.

The functional categorization of each superset was based on the known pathways from the Gene Ontology and KEGG databases. We evaluated the over-representation of a functional category within the member genes of a superset with the Fisher's exact test. Significant functional categories ($P < 0.05$ after Bonferroni correction) were used to annotate the functionality.

**Key driver analysis (KDA)**

We applied a previously developed key driver analysis (KDA) algorithm of gene-gene interaction networks [2-4] to the lipid-trait-associated supersets in order to identify the key regulatory genes. We used Bayesian gene networks due to their ability to capture detailed gene-gene relationships with causal implications [5]. In particular, we analyzed networks from several tissues, including adipose tissue, liver, blood, brain, kidney and muscle [6-14].

We defined a key driver as a gene that is connected to a large number of genes from a superset, compared to the expected number for a randomly selected gene within the Bayesian network. The algorithm searches first the neighbors of a target gene in the network, then calculates the ratio of neighbor genes that belong to a specific lipid-associated superset, and finally estimates the statistical significance by Fisher's exact test. As multiple networks were available for a number of tissues, we used two criteria to prioritize the five most important key genes. Firstly, we counted how many times a gene was a key driver in multiple datasets (denoted as $N$). The consistency across datasets was expressed as ($N$ - 0.99) to strongly favor key genes that could be identified in at least two datasets. The second criterion was based on the statistical significance of the key driver. In particular, the significance value was calculated as mean (log

***P***), where ***P*** denotes the Fisher's exact P-values from each of the networks. The final ranking was based on the product of the consistency and significance criteria. The SNP set enrichment and key driver analyses were performed using R.

**Supplemental References**

1. Zhong H, Yang X, Kaplan LM, Molony C, Schadt EE (2010) Integrating Pathway Analysis and Genetics of Gene Expression for Genome-wide Association Studies. American Journal of Human Genetics 86: 581-591.

2. Zhu J, Zhang B, Smith EN, Drees B, Brem RB, et al. (2008) Integrating large-scale functional genomic data to dissect the complexity of yeast regulatory networks. Nature Genetics 40: 854-861.

3. Wang IM, Zhang B, Yang X, Zhu J, Stepaniants S, et al. (2012) Systems analysis of eleven rodent disease models reveals an inflammatome signature and key drivers. Molecular Systems Biology 8.

4. Yang X, Zhang B, Molony C, Chudin E, Hao K, et al. (2010) Systematic genetic and genomic analysis of cytochrome P450 enzyme activities in human liver. Genome Research 20: 1020-1036.

5. Zhu J, Wiener MC, Zhang C, Fridman A, Minch E, et al. (2007) Increasing the power to detect causal associations by combining genotypic and expression data in segregating populations. Plos Computational Biology 3: 692-703.

6. Derry JMJ, Zhong H, Molony C, MacNeil D, Guhathakurta D, et al. (2010) Identification of Genes and Networks Driving Cardiovascular and Metabolic Phenotypes in a Mouse F2 Intercross. Plos One 5.

7. Emilsson V, Thorleifsson G, Zhang B, Leonardson AS, Zink F, et al. (2008) Genetics of gene expression and its effect on disease. Nature 452: 423-U422.

8. Fehrmann RSN, Jansen RC, Veldink JH, Westra HJ, Arends D, et al. (2011) Trans-eQTLs Reveal That Independent Genetic Variants Associated with a Complex Phenotype Converge on Intermediate Genes, with a Major Role for the HLA. Plos Genetics 7.

9. Greenawalt DM, Dobrin R, Chudin E, Hatoum IJ, Suver C, et al. (2011) A survey of the genetics of stomach, liver, and adipose gene expression from a morbidly obese cohort. Genome Research 21: 1008-1016.

10. Schadt EE, Molony C, Chudin E, Hao K, Yang X, et al. (2008) Mapping the genetic architecture of gene expression in human liver. Plos Biology 6: 1020-1032.

11. Wang SS, Schadt EE, Wang H, Wang XP, Ingram-Drake L, et al. (2007) Identification of pathways for atherosclerosis in mice - Integration of quantitative trait locus analysis and global gene expression data. Circulation Research 101: E11-E30.

12. Yang X, Schadt EE, Wang S, Wang H, Arnold AP, et al. (2006) Tissue-specific expression and regulation of sexually dimorphic genes in mice. Genome Research 16: 995-1004.

13. Tu ZD, Keller MP, Zhang CS, Rabaglia ME, Greenawalt DM, et al. (2012) Integrative Analysis of a Cross-Loci Regulation Network Identifies App as a Gene Regulating Insulin Secretion from Pancreatic Islets. Plos Genetics 8.

14. Nica AC, Parts L, Glass D, Nisbet J, Barrett A, et al. (2011) The Architecture of Gene Regulatory Variation across Multiple Human Tissues: The MuTHER Study. Plos Genetics 7.

15. Teslovich TM, Musunuru K, Smith AV, Edmondson AC, Stylianou IM, et al. (2010) Biological, clinical and population relevance of 95 loci for blood lipids. Nature 466: 707-713.

16. Ma L, Yang J, Runesha HB, Tanaka T, Ferrucci L, et al. (2010) Genome-wide association analysis of total cholesterol and high-density lipoprotein cholesterol levels using the Framingham Heart Study data. Bmc Medical Genetics 11.

17. Igl W, Johansson A, Wilson JF, Wild SH, Polasek O, et al. (2010) Modeling of Environmental Effects in Genome-Wide Association Studies Identifies SLC2A2 and HP as Novel Loci Influencing Serum Cholesterol Levels. Plos Genetics 6.

18. Aulchenko YS, Ripatti S, Lindqvist I, Boomsma D, Heid IM, et al. (2009) Loci influencing lipid levels and coronary heart disease risk in 16 European population cohorts. Nature Genetics 41: 47-55.

19. Tan AH, Sun JL, Xia N, Qin X, Hu YL, et al. (2012) A genome-wide association and gene-environment interaction study for serum triglycerides levels in a healthy Chinese male population. Human Molecular Genetics 21: 1658-1664.

20. Kim YJ, Go MJ, Hu C, Hong CB, Kim YK, et al. (2011) Large-scale genome-wide association studies in east Asians identify new genetic loci influencing metabolic traits. Nature Genetics 43: 990-U102.

21. Waterworth DM, Ricketts SL, Song KJ, Chen L, Zhao JH, et al. (2010) Genetic Variants Influencing Circulating Lipid Levels and Risk of Coronary Artery Disease. Arteriosclerosis Thrombosis and Vascular Biology 30: 2264-U2566.

22. Pollin TI, Damcott CM, Shen HQ, Ott SH, Shelton J, et al. (2008) A Null Mutation in Human APOC3 Confers a Favorable Plasma Lipid Profile and Apparent Cardioprotection. Science 322: 1702-1705.

23. Kathiresan S, Willer CJ, Peloso GM, Demissie S, Musunuru K, et al. (2009) Common variants at 30 loci contribute to polygenic dyslipidemia. Nature Genetics 41: 56-65.

24. Kathiresan S, Melander O, Guiducci C, Surti A, Burtt NP, et al. (2008) Six new loci associated with blood low-density lipoprotein cholesterol, high-density lipoprotein cholesterol or triglycerides in humans. Nature Genetics 40: 189-197.

25. Kooner JS, Chambers JC, Aguilar-Salinas CA, Hinds DA, Hyde CL, et al. (2008) Genome-wide scan identifies variation in MLXIPL associated with plasma triglycerides. Nature Genetics 40: 149-151.

26. Willer CJ, Sanna S, Jackson AU, Scuteri A, Bonnycastle LL, et al. (2008) Newly identified loci that influence lipid concentrations and risk of coronary artery disease. Nature Genetics 40: 161-169.

27. Saxena R, Voight BF, Lyssenko V, Burtt NP, de Bakker PIW, et al. (2007) Genome-wide association analysis identifies loci for type 2 diabetes and triglyceride levels. Science 316: 1331-1336.

28. Rasmussen-Torvik LJ, Pacheco JA, Wilke RA, Thompson WK, Ritchie MD, et al. (2012) High Density GWAS for LDL Cholesterol in African Americans Using Electronic Medical Records Reveals a Strong Protective Variant in APOE. Cts-Clinical and Translational Science 5: 394-399.

29. Trompet S, de Craen AJM, Postmus I, Ford I, Sattar N, et al. (2011) Replication of LDL GWAs hits in PROSPER/PHASE as validation for future (pharmaco)genetic analyses. Bmc Medical Genetics 12.

30. Shen HQ, Damcott CM, Rampersaud E, Pollin TI, Horenstein RB, et al. (2010) Familial Defective Apolipoprotein B-100 and Increased Low-Density Lipoprotein Cholesterol and Coronary Artery Calcification in the Old Order Amish. Archives of Internal Medicine 170: 1850-1855.

31. Sabatti C, Service SK, Hartikainen AL, Pouta A, Ripatti S, et al. (2009) Genome-wide association analysis of metabolic traits in a birth cohort from a founder population. Nature Genetics 41: 35-46.

32. Burkhardt R, Kenny EE, Lowe JK, Birkeland A, Josowitz R, et al. (2008) Common SNPs in HMGCR in Micronesians and Whites Associated With LDL-Cholesterol Levels Affect Alternative Splicing of Exon13. Arteriosclerosis Thrombosis and Vascular Biology 28: 2078-U2332.

33. Sandhu MS, Waterworth DM, Debenham SL, Wheeler E, Papadakis K, et al. (2008) LDL-cholesterol concentrations: a genome-wide association study. Lancet 371: 483-491.

34. Wallace C, Newhouse SJ, Braund P, Zhang F, Tobin M, et al. (2008) Genome-wide association study identifies genes for biomarkers of cardiovascular disease: Serum urate and dyslipiclemia. American Journal of Human Genetics 82: 139-149.

35. Weissglas-Volkov D, Aguilar-Salinas CA, Nikkola E, Deere KA, Cruz-Bautista I, et al. (2013) Genomic study in Mexicans identifies a new locus for triglycerides and refines European lipid loci. Journal of Medical Genetics 50: 298-308.

36. Wang K, Edmondson AC, Li M, Gao F, Qasim AN, et al. (2011) Pathway-Wide Association Study Implicates Multiple Sterol Transport and Metabolism Genes in HDL Cholesterol Regulation. Front Genet 2: 41.

37. Hiura Y, Shen CS, Kokubo Y, Okamura T, Morisaki T, et al. (2009) Identification of Genetic Markers Associated With High-Density Lipoprotein-Cholesterol by Genome-Wide Screening in a Japanese Population - The Suita Study. Circulation Journal 73: 1119-1126.

38. Ridker PM, Pare G, Parker AN, Zee RYL, Miletich JP, et al. (2009) Polymorphism in the CETP Gene Region, HDL Cholesterol, and Risk of Future Myocardial Infarction Genomewide Analysis Among 18 245 Initially Healthy Women From the Women's Genome Health Study. Circulation-Cardiovascular Genetics 2: 26-33.

39. Heid IM, Boes E, Muller M, Kollerits B, Lamina C, et al. (2008) Genome-Wide Association Analysis of High-Density Lipoprotein Cholesterol in the Population-Based KORA Study Sheds New Light on Intergenic Regions. Circulation-Cardiovascular Genetics 1: 10-U65.

40. Romanoski CE, Che N, Yin F, Mai N, Pouldar D, et al. (2011) Network for activation of human endothelial cells by oxidized phospholipids: a critical role of heme oxygenase 1. Circulation Research 109: e27-41.

41. Dimas AS, Deutsch S, Stranger BE, Montgomery SB, Borel C, et al. (2009) Common Regulatory Variation Impacts Gene Expression in a Cell Type-Dependent Manner. Science 325: 1246-1250.

42. Dixon AL, Liang L, Moffatt MF, Chen W, Heath S, et al. (2007) A genome-wide association study of global gene expression. Nature Genetics 39: 1202-1207.

43. Montgomery SB, Sammeth M, Gutierrez-Arcelus M, Lach RP, Ingle C, et al. (2010) Transcriptome genetics using second generation sequencing in a Caucasian population. Nature 464: 773-U151.

44. Stranger BE, Nica AC, Forrest MS, Dimas A, Bird CP, et al. (2007) Population genomics of human gene expression. Nature Genetics 39: 1217-1224.

45. Stranger BE, Montgomery SB, Dimas AS, Parts L, Stegle O, et al. (2012) Patterns of Cis Regulatory Variation in Diverse Human Populations. Plos Genetics 8: 272-284.

46. Duan S, Huang RS, Zhang W, Bleibel WK, Roe CA, et al. (2008) Genetic architecture of transcript-level variation in humans. American Journal of Human Genetics 82: 1101-1113.

47. Zeller T, Wild P, Szymczak S, Rotival M, Schillert A, et al. (2010) Genetics and Beyond - The Transcriptome of Human Monocytes and Disease Susceptibility. Plos One 5.
