## Supplemental Table S1 for "Gene Networks and Pathways for Plasma Lipid Traits via Multi-tissue Multi-omics Systems Analysis"

**Supplemental Table S1. Studies included in the positive control datasets for lipid traits.**

| Traits [a] | PubMed IDs | References | Reported Genes |
|---|---|---|---|
| TC | 20686565 | [15] | CELSR2, PSRC1, SORT1, APOE, APOC1, APOC2, LDLR, APOB, APOA1, APOC3, APOA4, APOA5, HMGCR, ABCG5, ABCG8, ANGPTL3, DOCK7, CSPG3, CILP2, PBX4, TRIB1, TIMD4, HAVCR1, ABCA1, GCKR, HP, HPR, DHX38, PCSK9, PPP1R3B, FADS1, FADS2, FADS3, ABO, LIPC, LIPG, HLA, TOP1, LPA, HNF1A, IRF2BP2, TOMM20, CETP, MOSC1, HNF4A, CYP7A1, BRAP, ST3GAL4, NPC1L1, TMEM57, LDLRAP1, C6orf106, MAFB, UBASH3B, FRK, FUT2, FLJ36070, GPAM, ERGIC3, DNAH11, PLEC1, NAT2, IDOL, TTC39B, RAF1, OSBPL7, RAB3GAP1, TRPS1, HFE, HIST1H4C, SPTY2D1, GVI1, EVI5, NCAN, TOMM40 |
| | 20370913 | [16] | |
| | 20066028 | [17] | |
| | 19060911 | [18] | |
| TG | 22171074 | [19] | APOA5, ALDH2, GCKR, TBL2, MLXIPL, LPL, ZNF259, APOA1, APOC3, APOA4, BUD13, APOB, DOCK7, ANGPTL3, BAZ1B, BCL7B, TRIB1, CILP2, ZNF101, AFF1, C5orf35, FADS1, APOE, APOC1, APOC2, CSPG3, PBX4, FADS2, FADS3, PLTP, HLA, GALNT2, NAT2, LIPC, CETP, JMJD1C, TIMD4, HAVCR1, KLHL8, FRMD5, ANKRD55, MAP3K1, COBLL1, LRP1, TYW1B, CCDC92, ZNF664, PINX1, XKR6, CAPN3, IRS1, CYP26A1, MSL2L1, CTF1, PLA2G6, DSCAML1, TOMM40, NCAN, AMAC1L2, ATG4C, LOC440069, MGC13125, C8orf35, SLC18A1, KIAA0999, LOC645044 |
| | 21909109 | [20] | |
| | 20864672 | [21] | |
| | 20686565 | [15] | |
| | 19074352 | [22] | |
| | 19060911 | [18] | |
| | 19060906 | [23] | |
| | 18193044 | [24] | |
| | 18193046 | [25] | |
| | 18193043 | [26] | |
| | 17463246 | [27] | |
| LDL | 23067351 | [28] | APOB, MYLIP, GMPR, PCSK9, CELSR2, HMGCR, TRIB1, BUD13, ZNF259, APOA5, APOA4, APOC3, APOA1, LDLR, SF4, CILP2, APOE, APOC1, APOC4, APOC2, PPP1R3B, PSRC1, SORT1, ABCG5, ABCG8, |
| | 21977987 | [29] | |
| | 21059979 | [30] | |

| | | | |
|---|---|---|---|
| | 20864672 | [21] | HP, HPR, DHX38, TIMD4, HAVCR1, CSPG3, PBX4, ABO, FADS1, FADS2, FADS3, TOP1, ANGPTL3, DOCK7, LPA, HNF1A, ST3GAL4, HLA, PLEC1, CETP, IRF2BP2, TOMM20, IDOL, NPC1L1, CBLN3, KIAA0323, MOSC1, TMEM57, LDLRAP1, HFE, HIST1H4C, DNAH11, BRAP, GPAM, FRK, CYP7A1, OSBPL7, MAFB, NCAN, TOMM40, CR1L, AR, B3GALT4, GCKR |
| | 20686565 | [15] | |
| | 19060911 | [18] | |
| | 19060906 | [23] | |
| | 19060910 | [31] | |
| | 18802019 | [32] | |
| | 18262040 | [33] | |
| | 18193044 | [24] | |
| | 18193043 | [26] | |
| | 18179892 | [34] | |
| HDL | 23505323 | [35] | SLC12A3, NLRC5, HERPUD1, CETP, TSPAN16, SPC24, RAB3D, LOC55908, KANK2, DOCK6, PPP1R3B, NIPSNAP3A, NIPSNAP3B, ABCA1, MYL2, C12orf51, OAS3, LPL, PVRL2, TOMM40, APOE, FADS1, TTC39B, GALNT2, APOB, ZNF259, APOA5, APOA4, APOC3, APOA1, MYO1H, KCTD10, UBE3B, MMAB, MVK, LIPC, GFOD2, LCAT, LIPG, FADS2, FADS3, PLTP, APOC1, APOC2, TRIB1, LRP4, NR1H3, LILRA3, LILRB2, HNF4A, KLF14, SCARB1, STARD3, CMIP, TRPS1, SLC39A8, ABCA8, COBLL1, CCDC92, ZNF664, ZNF648, MACF1, PABPC4, MLXIPL, IRS1, C6orf106, PGS1, RPS3A, MC4R, SBNO1, LACTB, UBE2L3, LRP1, CITED2, UBASH3B, LPA, ANGPTL4, PDE3A, ADM, AMPD3, ARL15, NUP93, MADD, FOLH1, CTCF, PRMT8, ACAA2 |
| | 21909109 | [20] | |
| | 22303337 | [36] | |
| | 20864672 | [21] | |
| | 20686565 | [15] | |
| | 19359809 | [37] | |
| | 20031564 | [38] | |
| | 19060911 | [18] | |
| | 19060906 | [23] | |
| | 19060910 | [31] | |
| | 20031538 | [39] | |
| | 18193044 | [24] | |
| | 18193043 | [26] | |

Traits [a]: TG: Triglycerides; LDL: Low-Density Lipoprotein Cholesterol; HDL: High-Density Lipoprotein Cholesterol.
