## Supplemental Table S2 for "Gene Networks and Pathways for Plasma Lipid Traits via Multi-tissue Multi-omics Systems Analysis"

**Supplemental Table S2. Studies that produced the gene co-expression modules and Bayesian network models of gene-gene interactions.**

| Tissues | Data sources | References |
|---|---|---|
| Human gene expression resources and gene interaction networks | | |
| Aortic endothelial cells | 149 heart transplant donors | [40] |
| Adipose tissue and blood | 1,675 individuals from two Icelandic cohorts | [7] |
| Blood | 1,469 unrelated individuals | [8] |
| Fibroblasts, lymphoblastoid cells and T cells | Umbilical cords of 85 Western European individuals | [41] |
| Liver | 427 individuals | [10] |
| Liver and adipose tissue | 1,008 obese patients | [9] |
| Lymphoblastoid cells | 400 children of families with a proband with asthma | [42] |
| | 60 HapMap participants of European descent | [43] |
| | 270 HapMap participants | [44] |
| | 726 HapMap3 participants | [45] |
| | 30 European and 30 Yoruba HapMap participants | [46] |
| Lymphoblastoid cells, skin and adipose tissue | 150 female twins | [14] |
| Monocytes | 1,490 unrelated individuals | [47] |
| Mouse gene expression resources and gene interaction networks | | |
| Liver, adipose tissue, kidney, heart, muscle, brain | C57BL/6J x A/J cross | [6] |
| Liver, adipose tissue, muscle, brain | C57BL/6J x C3H ApoE -/- | [11,12] |
| Liver, adipose tissue, muscle | C57BL/6J x C3H wildtype | [10] |
| Islet cells, adipose tissue, liver, muscle, hypothalamus | C57BL/6J x BTBR $Lep^{ob}$ | [13] |
