## Supplemental Table S3 for "Gene Networks and Pathways for Plasma Lipid Traits via Multi-tissue Multi-omics Systems Analysis"

### Supplemental Table S3. Primer pairs for RT qPCR

| *Genes* | *Forward (5'-3')* | *Reverse (5'-3')* |
|---|---|---|
| *F2* | *TTCCCGAACCAGATATGAGC* | *CTGTCTGCTTGTCTGGCAAA* |
| *F2 siRNA-1* | *GGAACAGCUUACCAGCCAATT* | *UUGGCUGGUAAGCUGUUCCTT* |
| *F2 siRNA-2 (chosen)* | *GCGAUAUCGCUCUGCUCAATT* | *UUGAGCAGAGCGAUAUCGCTT* |
| *F2 siRNA-3* | *GCUCCGGGCUGGUUAUAAATT* | *UUUAUAACCAGCCCGGAGCTT* |
| *Sc siRNA* | *UUCUCCGAACGUGUCACGUTT* | *ACGUGACACGUUCGGAGAATT* |
| *Abcb11* | *GGACAATGATGTGCTTGTGG* | *CACACAAAGCCCCTACCAGT* |
| *Apoa4* | *GTCACCTTCCAACGTGGAGT* | *GATGTTTCCCGGAGTCAGAA* |
| *Apoa5* | *GAGTCGAGTGCTGCACCATA* | *ACGTGTGAGTTTGTGGGACA* |
| *Apof* | *GAAAGGATGCCACCACAGAT* | *CCACCCATCCATCCATCTAC* |
| *Fabp1* | *GGAAGGACATCAAGGGGGTG* | *TCACCTTCCAGCTTGACGAC* |
| *Gc* | *ACTCAGAGTCCCCTGCTGAA* | *GGTGTTGGGTGTTTTTGTCC* |
| *Hrg* | *ATGGTCCACCACATGGACAC* | *AGTGGAGGGAGTCGGTAGAC* |
| *Lipc* | *GACTGGATCTCCCTGGCATA* | *AGGTGAACTTTGCTCCGAGA* |
| *Plg* | *GCTGCCTGTGATTGAGAACA* | *CTCGAAGCAAACCAGAGGTC* |
| *Proc* | *GGACCTGGACATCAAGGAGA* | *GCAGATGGGCACTATGGTTT* |
| *Snrbp2* | *AAGAGATCCCTGTATGCCCTT* | *GTGGATGAACCCAGTTCCTTAAA* |
| *Gpt* | *TCCAGGCTTCAAGGAATGGAC* | *CAAGGCACGTTGCACGATG* |
| *Itga6* | *TGCAGAGGGCGAACAGAAC* | *GCACACGTCACCACTTTGC* |
| *Spry44* | *GCAGCGTCCCTGTGAATCC* | *TCTGGTCAATGGGTAAGATGGT* |
| *Lep* | *GAGACCCCTGTGTCGGTTC* | *CTGCGTGTGTGAAATGTCATTG* |
| *Pparg* | *CCATTCTGGCCCACCAAC* | *AATGCGAGTGGTCTTCCATCA* |
| *Cebpa* | *GCGGGCAAAGCCAAGAA* | *GCGTTCCCGCCGTACC* |
| *Srebp1* | *CTCAGCAGCCACCATCTAGCCT* | *GCTGATGCCTGCAGTCTTCACG* |
| *Fasn* | *CTG AGATCCCAGCACTTCTTGA* | *GCCTCCGAAGCCAAATGAG* |
| *Adipoq* | *GATGGCACTCCTGGAGAGAA* | *TCTCCAGGCTCTCCTTTCCT* |
| *Lipe* | *ACAGTGCAGGTGGGAATCTC* | *GCCTAGTGCCTTCTGGTCT* |
| *Cd36* | *GCAGGTCTATCTACGCTGTG* | *GGTTGTCTGGATTCTGGAGG* |
| *Fabp4* | *TGAAATCACCGCAGACGACA* | *ATAACACATTCCACCACCAGC* |
| *Beta actin* | *GCAGGAGTACGATGAGTCCG* | *ACGCAGCTCAGTAACAGTCC* |
