## Supplemental Figure S1 for "Gene Networks and Pathways for Plasma Lipid Traits via Multi-tissue Multi-omics Systems Analysis"

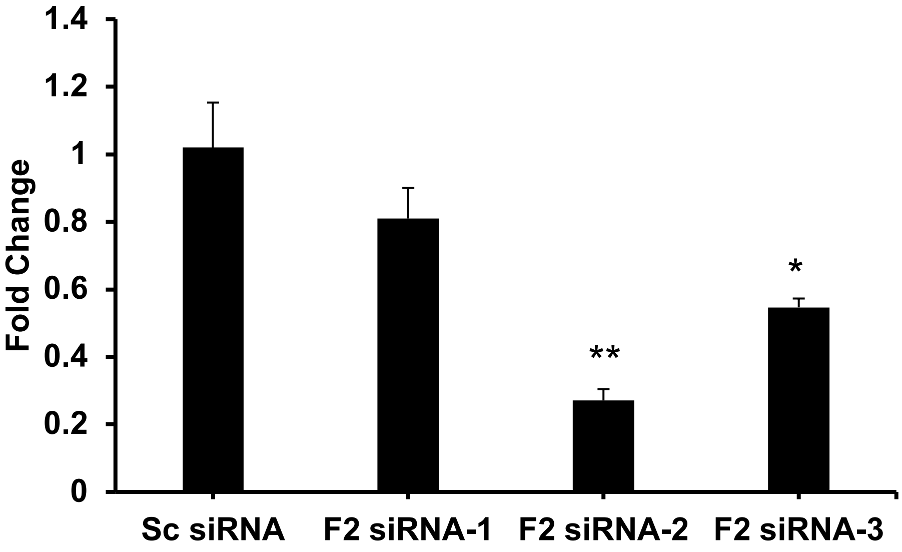


**Supplemental Figure S1. Gene knockdown efficiencies of three *F2* siRNAs.** Three *F2* siRNAs were tested to select for the most efficient knockdown of the target gene. 3T3-L1 adipocytes were transfected with *F2* siRNA at day 7 of differentiation (D7). Scrambled (Sc) siRNA was used as the control for normalization. After 48 h, *F2* gene expression was analyzed by real-time qPCR. *Beta actin* was used as a housekeeping gene. Result represents the mean ± s.e.m. n = 3/group. Statistical significance was determined by Student's t-test between each *F2* siRNA and the Sc siRNA (*$p < 0.05$ and **$p < 0.01$).
