## Supplemental Figure S2 for "Gene Networks and Pathways for Plasma Lipid Traits via Multi-tissue Multi-omics Systems Analysis"

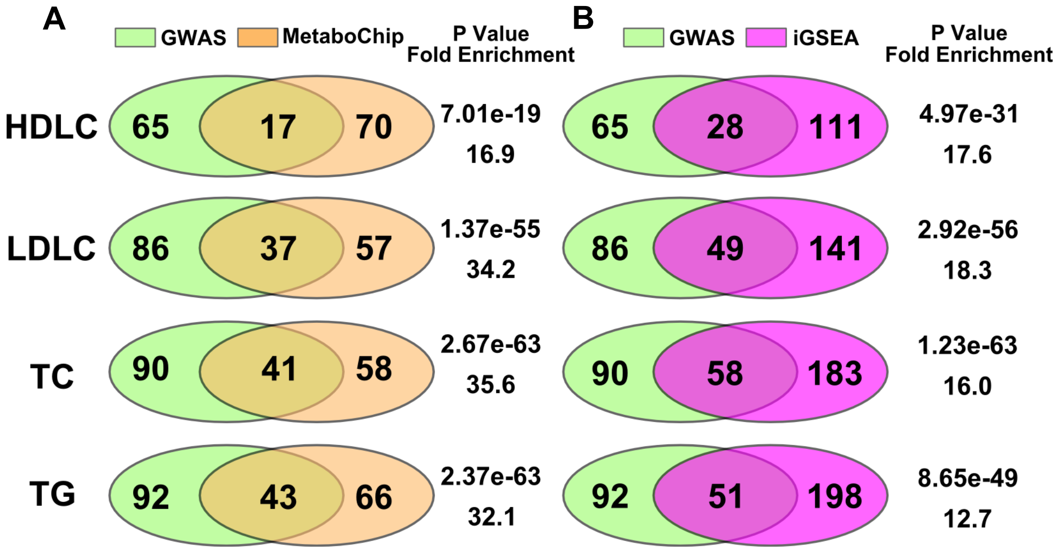


**Supplemental Figure S2. Validation of MSEA results from GLGC GWAS using independent genetic association data from MetaboChip and a different method iGSEA. A)** Venn diagram of the convergent pathways between GLGC GWAS and MetaboChip dataset using the same MSEA method. **B)** Venn diagram of the convergent pathways between MSEA and iGSEA for the same GLGC GWAS dataset. Fisher exact test was applied to evaluate the overlap in the pathways detected using different datasets or using different methods, with 4532 pathways in total.
