## Supplemental Figure S3 for "Gene Networks and Pathways for Plasma Lipid Traits via Multi-tissue Multi-omics Systems Analysis"

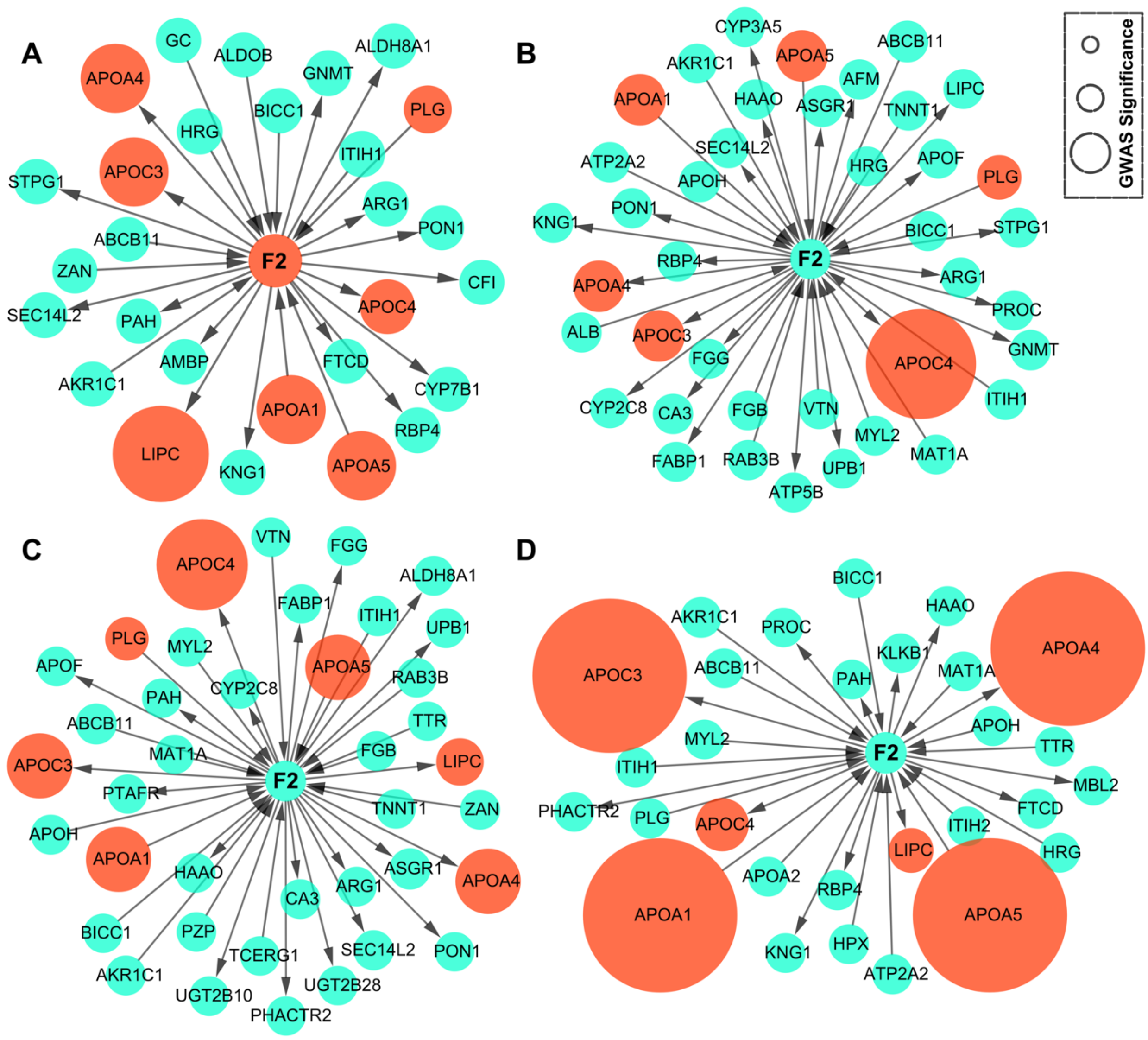


**Supplemental Figure S3. GWAS genes in Neighboring genes of Gene *F2* in human Bayesian networks.** Panel **(A-D)** represent GWAS susceptibility genes around gene *F2* for HDL, LDL, TC, and TG respectively. The interactions come from a combined Bayesian network from different human tissues, including adipose, liver, blood, kidney, muscle, and brain. The node size corresponds to the GWAS significance.
